# ATAD3A structurally links mtDNA replication and mitochondrial fission

**DOI:** 10.64898/2026.06.23.733767

**Authors:** Nitish Dua, Boyuan Ma, Samantha Oviedo, Hamidreza Rahmani, Tumara Boyd, Donghyun Park, R. Luke Wiseman, Danielle A. Grotjahn

**Affiliations:** Department of Integrative Structural and Computational Biology, The Scripps Research Institute, La Jolla, CA 92037, USA; Department of Molecular and Cellular Biology, The Scripps Research Institute, La Jolla, CA 92037, USA

## Abstract

Mitochondrial function depends on the maintenance of its genome, and disruptions in copy number and distribution are hallmarks of mitochondrial disorders. Mitochondrial DNA (mtDNA) replication is spatially and temporally linked to mitochondrial division (i.e., fission). However, the signal that coordinates these two events, which are physically separated by the barrier of two mitochondrial membranes, remains unknown. To gain insight into this coordination, we employed correlative cryo-electron tomography (cryo-ET) to analyze the microenvironment surrounding replicating nucleoids. Mitochondrial regions containing replicating mtDNA exhibit a unique membrane architecture defined by the presence of clustered, membrane-spanning tethers that traverse the inner membrane space. Using a combination of superresolution microscopy and genetically encoded cryo-ET tagging technology, we identify these tethers as the AAA+ ATPase ATAD3A. We further show that ATAD3A knockdown reduces recruitment of the mitochondrial fission machinery, whereas overexpression promotes its recruitment and subsequent fission. Our work suggests that ATAD3A forms nanoscale linkages that coordinate these two distinct processes, revealing a new structural paradigm for organellar communication across distinct membrane-defined environments.

**Highlights:**

- Replicating mitochondrial DNA (mtDNA) nucleoids are surrounded by a distinct membrane microenvironment.
- ATAD3A forms membrane-spanning tethers enriched at replicating mtDNA sites.
- ATAD3A enrichment is necessary and sufficient to recruit mitochondrial fission machinery and induce fission at mtDNA replication sites.
- ATAD3A structurally couples mtDNA replication state to mitochondrial fission across distinct organellar subcompartments.

## Introduction

The human mitochondrial genome is indispensable for cellular function as it encodes 13 proteins essential for electron transport chain function, along with tRNAs and rRNAs required for translation Garrido et al. (2003); Miyakawa (2017). Mitochondrial DNA (mtDNA) is packaged into functional units in the form of nucleo-protein complexes, termed ‘nucleoids’, which are distributed throughout the mitochondrial network. The maintenance of mitochondrial DNA (mtDNA) copy number and distribution is vital for proper mitochondrial function, and perturbations in mtDNA copy number are a hallmark of mitochondrial disorders Filograna et al. (2021). Hence, it is important to understand how cells maintain these genomes to preserve cellular function Craven et al. (2017); Tuppen et al. (2010); Taylor and Turnbull (2005)

The replication of mtDNA and mitochondrial division (i.e., fission) are spatially and temporally linked through the recruitment of cytosolic factors such as dynamin-related protein 1 (DRP1) and the endoplasmic reticulum (ER) Kleele et al. (2021); Lewis et al. (2016); Newman et al. (2024); Friedman et al. (2011). The coordination between the two events not only ensures faithful replication but also facilitates segregation of the newly replicated copies and their distribution within the network. This is evident from studies showing that enlarged mtDNA nucleoids accumulate in cells when mitochondrial fission is inhibited Ishihara et al. (2022); Ban-Ishihara et al. (2013). Despite this established spatiotemporal coordination, the mechanism generating cellular communication between ‘internal’ mtDNA nucleoids and ‘external’ fission factors in the cytosol remains unclear.

To address the coupling between mtDNA replication and fission, we developed a cryo-correlative light and electron tomography (cryo-ET) pipeline to define the structural changes that occur in the local microenvironment of the mtDNA nucleoid during replication. We hypothesized that mtDNA replication could influence its local environment, thereby inducing characteristic changes and enabling communication with the cytosolic fission machinery. We find that mtDNA nucleoids are uniquely harbored in mitochondrial subdomains of uniform inner-to-outer membrane distances, which are enriched in ‘tether’ densities that span these two membranes. Using a recently described genetically encoded cryo-ET tagging technology Fung et al. (2023), we identify these intermembrane tethers as the AAA+ ATPase containing protein 3A, ATAD3A, and provide evidence for a role for these tethers in recruiting fission components. Collectively, our findings reveal that ATAD3A forms nanoscale intermembrane linkages that spatially coordinate mtDNA replication and fission, redefining the structural basis for linking mitochondrial dynamics and genome maintenance across distinct organellar compartments.

## Results

### mtDNA replication is associated with a unique mitochondrial membrane microenvironment

To visualize the local subcellular environment of replicating nucleoids, we transfected cells with a plasmid containing mScarlet-POLG2 as a marker to specifically label replicating nucleoids Lewis et al. (2016) and FACS-sorted transfected cells directly on EM grids to enrich for mScarlet-POLG2-expressing cells (Fig. 1A, Fig. S1A,B). We then treated cells with a short pulse of SYBRgold dye to stain the total mtDNA population prior to cell vitrification via plunge-freezing. Using cryo-fluorescence microscopy, we identified and then subsequently targeted fluorescent cells for cryo-focused ion beam (cryo-FIB) milling to generate thin cellular sections (i.e., lamellae). We imaged each lamella using an in-chamber cryo-fluorescence microscopy to determine the precise position of SYBRgold and mScarlet-POLG2 fluorescence puncta (Fig. 1B, C). We then acquired a high-magnification transmission electron microscopy (TEM) montage of each lamella and targeted mitochondria-containing regions for tilt series collection (Fig. 1D). The post cryo-FIB-milling fluorescence image was overlaid on the TEM overview to correlate to classify all mitochondria within our tomographic data into three different categories (i.e., mtDNA “states”): no fluorescence signal (no DNA, *n* = 43 mitochondria), only SYBRgold positive (non-replicating mtDNA, *n* = 28 mitochondria), or both SYBRgold and mScarlet-POLG2 positive (replicating mtDNA, *n* = 26 mitochondria). (Fig. S1C). Mitochondria that did not contain any fluorescence signal would represent parts of the mitochondria network that do not contain an mtDNA nucleoid, or, in some cases, could also represent mitochondria in which the fluorescent signal was milled away during cryo-FIB milling.

**Figure 1.**
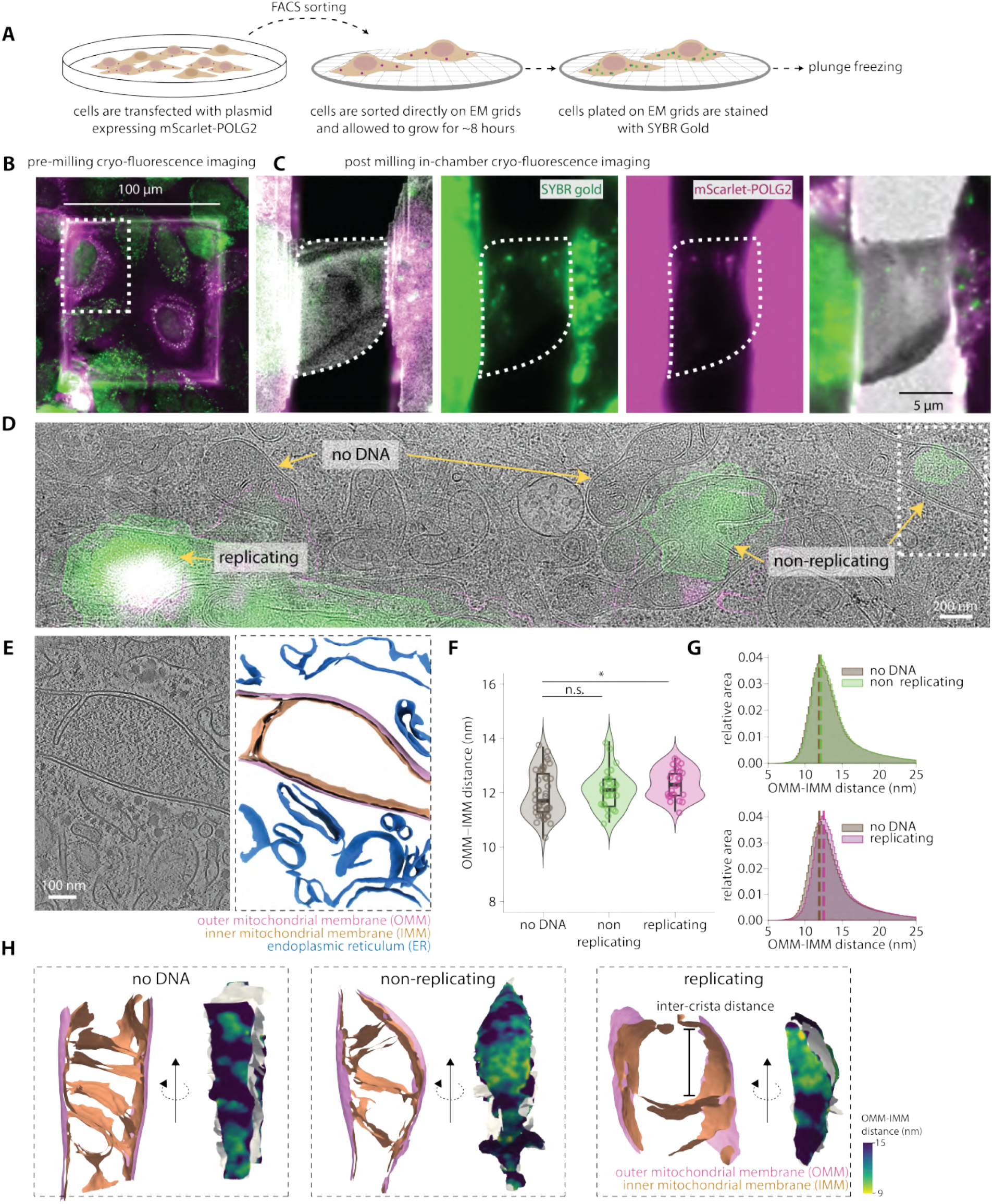
mtDNA replication is associated with a unique membrane microenvironment. (**A**) Schematic showing workflow for labelling HeLa cells with mScarlet-POLG2 and SYBRGold prior to vitrification. (**B**) Representative images showing part of the EM grid imaged on a cryo-fluorescence microscope. (**C**) Images acquired post cryo-FIB milling of the cell marked by a white dotted rectangle shown in B. Fluorescence images are acquired on an in-chamber cryo-fluorescence microscope and correlated with the SEM image. The fluorescence image is also correlated with a TEM overview of the lamella (right). (**D**) The TEM overview is correlated with the post-milling cryo-fluorescence image shown in C. White rectangles with dotted lines mark a representative region where tilt series were acquired. (**E**) Representative z-slice from a denoised 3D tomogram and membrane segmentations with outer mitochondrial membrane (OMM), inner mitochondrial membrane (IMM), and endoplasmic reticulum (ER) membrane annotated. (**F**) Peak histogram values from each mitochondrion for OMM-IMM distance are plotted as a violin plot for all three classes. Each dot represents peak value for the corresponding mitochondrion. (number of mitochondria = 43 no DNA; 28 non-replicating, 26 replicating). (**G**) Combined histogram of OMM–IMM distances for all three classes, dashed vertical lines correspond to peak histogram values of pooled data. (number of mitochondria = 43 no DNA; 28 non-replicating, 26 replicating) (**H**) Membrane surface reconstructions for representative examples from three classes of mitochondria are shown with per triangle measurements of OMM-IMM distance mapped on the surface as a color gradient. Statistical significance in F is calculated using Mann-Whitney U test (\*\*\**p<* 0.001, \*\**p<* 0.01, \**p<* 0.05).

We segmented membranes from the resulting three-dimensional (3D) tomograms (Fig. 1E) and used our ‘surface morphometrics’ pipeline Barad et al. (2023) to compare the membrane ultrastructure of mitochondria across all three mtDNA states. Mitochondria containing mtDNA displayed larger inter-crista distance as expected Liu et al. (2022); Stephan et al. (2019) (Fig. S1D and Fig. S2). We observed a significant reduction in the variability of outer-to-inner (OMM-IMM) distances of mitochondria containing replicating mtDNA relative to mitochondria with non-replicating mtDNA or no mtDNA, with a median distance of ∼12.5 nm between the two membranes (Fig. 1F–H and Fig. S2). To ensure that our observations were not skewed by differences in the number of cristae between categories, we quantified OMM-IMM distance only for the inner boundary membrane (IBM) regions of the mitochondria and observed similar distributions (Fig. S1E,F). Additionally, we found that mitochondria with replicating mtDNA exhibited a higher frequency of larger patches of membrane (∼300– 400 nm) with an OMM-IMM distance of 12–13 nm relative to the other two correlation groups (Fig. S1G,H). Collectively, these results show that mitochondria harboring mtDNA have localized regions of uniform OMM-IMM distances, suggesting that local membrane remodeling is associated with mtDNA replication.

### Mitochondria with replicating mtDNA contain clustered, membrane-spanning tethers

We hypothesized that the uniform OMM-IMM distances observed in mitochondria with replicating mtDNA were mediated by proteins that physically link opposing bilayers. Consistent with this, we observed “tether”-like densities in the intermembrane space regions exhibiting ∼ 12 nm OMM-IMM distances (Fig. 2A–C). We performed blinded, manual two-point particle picking of these tethers (Fig. 2C) across all tomograms in our three correlation groups, followed by subtomogram averaging, which resulted in a strong 12 nm density between the two appressed membranes (Fig. 2D). As a control, we performed subtomogram averaging on an equal number of randomly selected patches of the membrane. This resulted in a structure with a clear membrane signal but no signal within the inner membrane space, suggesting that the bridging density we observe in our tomograms is a specific signal rather than random noise (Fig. S3A).

**Figure 2.**
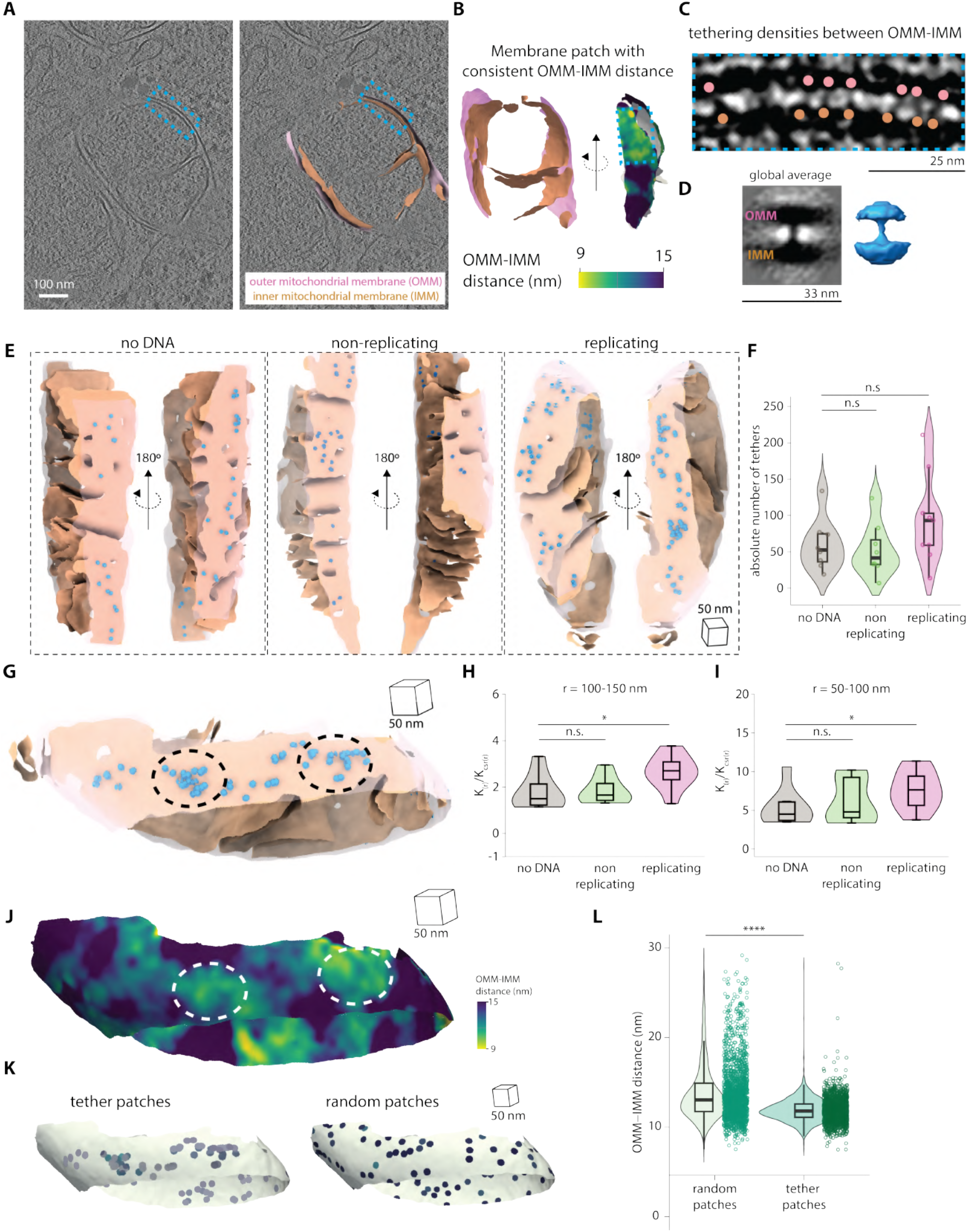
Regions of uniform OMM-IMM distance contain tether-like densities that are clustered at replicating mtDNA nucleoids. (**A**) Representative z-slice from a 3D tomogram in the “replicating” category with membrane surfaces overlayed. The blue dotted rectangle shows part of the membrane shown in B and C. (**B**) The blue dotted lines mark part of the membranes showing a uniform OMM-IMM distance patch. (**C**) Zoomed-in view of the 3D tomogram showing tether-like densities spanning OMM-IMM. Dots represent the two end points of a tether on both the OMM (salmon) and the IMM (orange), where dipole picking for sub-tomogram extraction and averaging was performed. (**D**) Subtomogram global average from 1,861 particles shows density between OMM and IMM. (**E**) The global average from D (blue) is mapped onto the mitochondrial surfaces of the respective three classes of mitochondria. (**F**) The number of tethers in each category of mitochondria is plotted as a violin plot. Each data point represents one mitochondrion. (*n* = 8 no DNA; 8 non-replicating, 9 replicating) (**G**) Representative example of mitochondrion from the “replicating” category with dotted black oval marking clusters of tethers. (**H, I**) Quantification of the maximum value of *K*(*r*)*/K*_CSR_(*r*) for each tomogram at the indicated radius intervals in each category. (*n* = 8 no DNA; 8 non-replicating, 9 replicating) (**J**) Per-triangle measurements of OMM-IMM distance mapped on the surface for mitochondria shown in G, showing how the same regions of clustered tethers overlap with regions of consistent OMM-IMM distances (white dashed oval). (**K**) Extracted membrane patches of diameter 10 nm at sites possessing a tether, or random patches. (**L**) OMM-IMM distance is plotted as a violin plot for random and tether containing patches of radius 10 nm. Each data point represents the median OMM-IMM distance for a patch. (1,861 tether patches and 1,861 random patches from *n* (mitochondria) =8 no DNA; 8 non-replicating, 9 replicating). Statistical significance in F, H, I and L is calculated using Mann-Whitney U test (\*\*\*\**p<* 0.0001, \*\*\**p<* 0.001, \*\**p<* 0.01, \**p<* 0.05).

Mapping the location of every tether onto the mitochondrial membrane surface showed an increase in the total number of tethers in mitochondria with replicating mtDNA (Fig. 2E, F). A similar trend was observed when the number of tethers was normalized to outer-membrane surface area (Fig. S3B). We also observed a significant increase in the propensity of these tethers to cluster in mitochondria with replicating mtDNA (Fig. 2G), as quantified by Ripley’s K function (see Methods). This clustering occurred across large-scale distances (50–100 nm and 100–150 nm) in mitochondria containing replicating mtDNA (Fig. 2H, I), with no significant clustering observed for radii lower than 50 nm or more than 150 nm (Fig. S3C). This suggests that the presence of large clusters of tethers spanning extensive membrane patches is a unique feature of mitochondria with replicating mtDNA.

We hypothesized that individual tethers may exhibit a uniform local distance across the intermembrane space, and that clustering of these tethers may facilitate the large patches of uniform OMM-IMM distance observed in mitochondria with replicating mtDNA (Fig. 2G, J). To test this hypothesis, we computationally extracted a local patch of OMM (radius = 10 nm), centred on the tether and measured the OMM-IMM distance in this region. We found that the OMM-IMM distance at tether-associated patches exhibited a very tight distribution around 12 nm, whereas patches generated randomly across the surface showed a much wider spread and greater variability (Fig. 2K, L). These distributions were the same across all three categories of mitochondria (Fig. S3D). This suggests that the tethers maintain a consistent OMM-IMM distance regardless of where they are in the membrane, but that the increase in the abundance and clustering of these tethers leads to local remodeling, resulting in larger patches of uniform OMM-IMM distances in mitochondria containing replicating mtDNA.

### ATAD3A is enriched at replicating mtDNA nucleoids

We hypothesized that these tethers could represent the AAA+ ATPase family protein ATAD3A, since it is the only known protein in mammals shown to span both the inner and outer mitochondrial membranes and is involved in mtDNA nucleoid dynamics Ishihara et al. (2022); Gilquin et al. (2010); Arguello et al. (2021); Cooper et al. (2017); Zhao et al. (2019); Harel et al. (2016); Peralta et al. (2018). Additionally, ATAD3A can bind to DRP1 via its N-terminus and, independently, to the D-loop of mtDNA Zhao et al. (2019); He et al. (2007), making it a potential candidate for linking mtDNA and mitochondrial fission. Since other members of the AAA+ ATPase family have been shown to form hexamers Puchades et al. (2020), we performed AlphaFold modelling Abramson et al. (2024) for ATAD3A isoform 1 and isoform 2 hexamers independently, resulting in structure predictions for both isoforms that form a stalk-like structure, with the C-terminus ATPase domains forming the canonical ‘ring’ structure on one side (Fig. 3A). We generated a model of the AlphaFold structure with membrane densities (CHARMM Jo et al. (2008)) by positioning the IMM at the predicted transmembrane region of ATAD3A close to the ATPase domains and by positioning OMM close to the N-terminus region (Fig. 3B). Using this model, we simulated a 3D tomogram Purnell et al. (2023) and visualized the stalk-like connection between the two membranes, measuring ∼12 nm in length, consistent with the distance observed in the sub-tomogram average (Fig. 3B and Fig. S4A). This suggests that ATAD3A hexamer structural prediction models match the dimensions and features of the tethers we observe spanning the intermembrane space in mitochondria within our cellular cryo-ET data.

**Figure 3.**
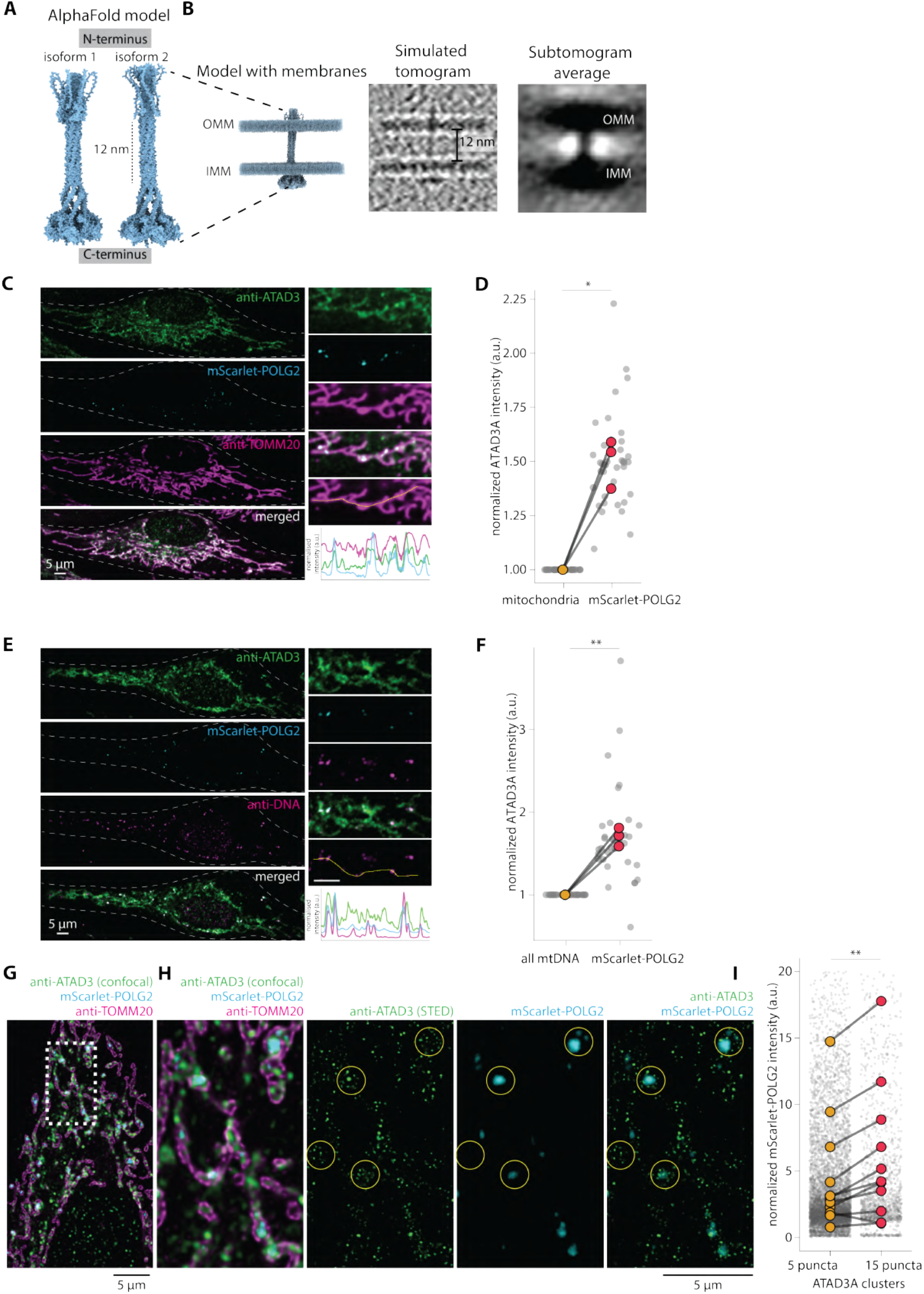
ATAD3A is enriched at replicating mtDNA nucleoids. (**A**) An AlphaFold model of ATAD3A isoform 1 hexamer and isoform 2 hexamer. (**B**) The model generated using ATAD3A isoform 1 AlphaFold prediction and OMM and IMM PDB models from CHARMM *(left)*, simulated tomogram of the ATAD3A hexamer and membrane model *(middle)*, along with the sub-tomogram average of the tethers *(right)*. (**C**) Representative confocal images of HeLa cells expressing mScarlet-POLG2 and stained with anti-TOMM20 and anti-ATAD3A antibodies. The inset shows part of the cell with a line profile. (**D**) Fold change in normalized ATAD3A intensity over mScarlet-POLG2 puncta, as compared to the entire mitochondrial network within the same cell, is plotted. Each grey data point represents one cell, and the lines connect the means across 3 biological replicates. (*n* = 74 cells, from 3 replicates) (**E**) Representative confocal images of HeLa cells expressing mScarlet-POLG2 and stained with anti-DNA and anti-ATAD3A antibodies. The inset shows part of the cell with a line profile. (**F**) Fold change in normalized ATAD3A intensity over mScarlet-POLG2 puncta, as compared to all mtDNA within the same cell, is plotted. Each grey data point represents one cell, and lines connect means for 3 different biological replicates (*n* = 68 cells, from 3 replicates). (**G**) Representative confocal image of HeLa cells expressing mScarlet-POLG2 and stained with anti-TOMM20 and anti-ATAD3A antibodies. (**H**) Inset from G shows part of the cell with STED resolved ATAD3A clusters, and yellow lines mark the identified clusters of ≥15 puncta within 0.5 *µ*m. (**I**) Normalized mScarlet-POLG2 intensity over identified ATAD3A clusters with ≤5 or ≥15 puncta within 0.5 *µ*m. Each grey dot represents one identified cluster, and lines connect means of each cell. (*n* = 12 cells, from 2 replicates). Statistical significance in D, F, and I is calculated using a paired *t* -test (\*\*\*\**p<* 0.0001, \*\*\**p<* 0.001, \*\**p<* 0.01, \**p<* 0.05).

To test whether ATAD3A exhibits localization signatures similar to those of the tethers, we used immunofluorescence to localize ATAD3A in cells alongside mtDNA. Consistent with previous studies, we found that ATAD3A was distributed throughout the mitochondrial network Arguello et al. (2021); He et al. (2007); however, there was a significant enrichment of ATAD3A signal around sites of replicating mtDNA relative to the entire mitochondrial network and all mtDNA puncta (Fig. 3C–F). To further characterize this enrichment, we used stimulated emission depletion (STED) nanoscopy, which revealed that individual puncta visualized by diffraction-limited confocal microscopy comprised multiple ATAD3A signal puncta. (Fig. 3G, H). We used an unbiased approach to identify clusters of ≥5 puncta or ≥15 puncta within a fixed radius of 0.5 *µ*m of STED-resolved ATAD3A and found that, while ≥ 5 puncta clusters were identified across the entire mitochondrial network, ≥15 clusters were restricted to parts of mitochondria positive for mScarlet-POLG2 (Fig. 3H, I, and Fig. S4B). Collectively, these results demonstrate that ATAD3A clusters specifically around replicating mtDNA in a manner similar to the tethers we observe in our cryo-ET, suggesting that ATAD3A enrichment is a unique feature of the local environment surrounding replicating mtDNA.

### ATAD3A assembles as an intermembrane space-spanning tether in mitochondria

We next sought to confirm that the intermembrane space-spanning tethers we observe in mitochondria with replicating mtDNA in our correlative cryo-ET data represent ATAD3A, which we observe as enriched clusters in our immunofluorescence experiments. Historically, the localization of specific proteins in cellular cryoET data has been challenging; however, the recent development of genetically encoded tags has enabled the subcellular localization of some protein targets, including those localized to the OMM Fung et al. (2023). We developed a system using the genetically encoded multimeric (GEM) cryo-ET tag, which spontaneously forms a virus-like icosahedral particle in cells and can be attached to a GFP-tagged variant of a protein of interest through rapalog coupling Fung et al. (2023). We generated an N-terminal GFP-tagged version of ATAD3A isoform 1, which, based on previous work and our AlphaFold model, we predicted would expose the GFP coupled to the N-terminus of ATAD3A in the cytosol and make it available for GEM tag binding (Fig. 4A, B). We confirmed the co-localization of GFP-ATAD3A with mitochondria via confocal microscopy and also found that the GFP-ATAD3A signal overlapped with STED-resolved TOMM20 puncta (Fig. 4A). We co-expressed GFP-ATAD3A and GEM donor plasmid in HeLa cells and treated with rapalog (i.e., coupled heterodimerization between GEM tag and GFP-ATAD3A) prior to processing through our correlative cryo-ET pipeline (as described earlier in Fig. 1, see Methods). As a control, cells expressing both constructs in the absence of rapalog treatment (i.e., uncoupled) were prepared for cryo-ET imaging (Fig. 4B) (see Methods).

**Figure 4.**
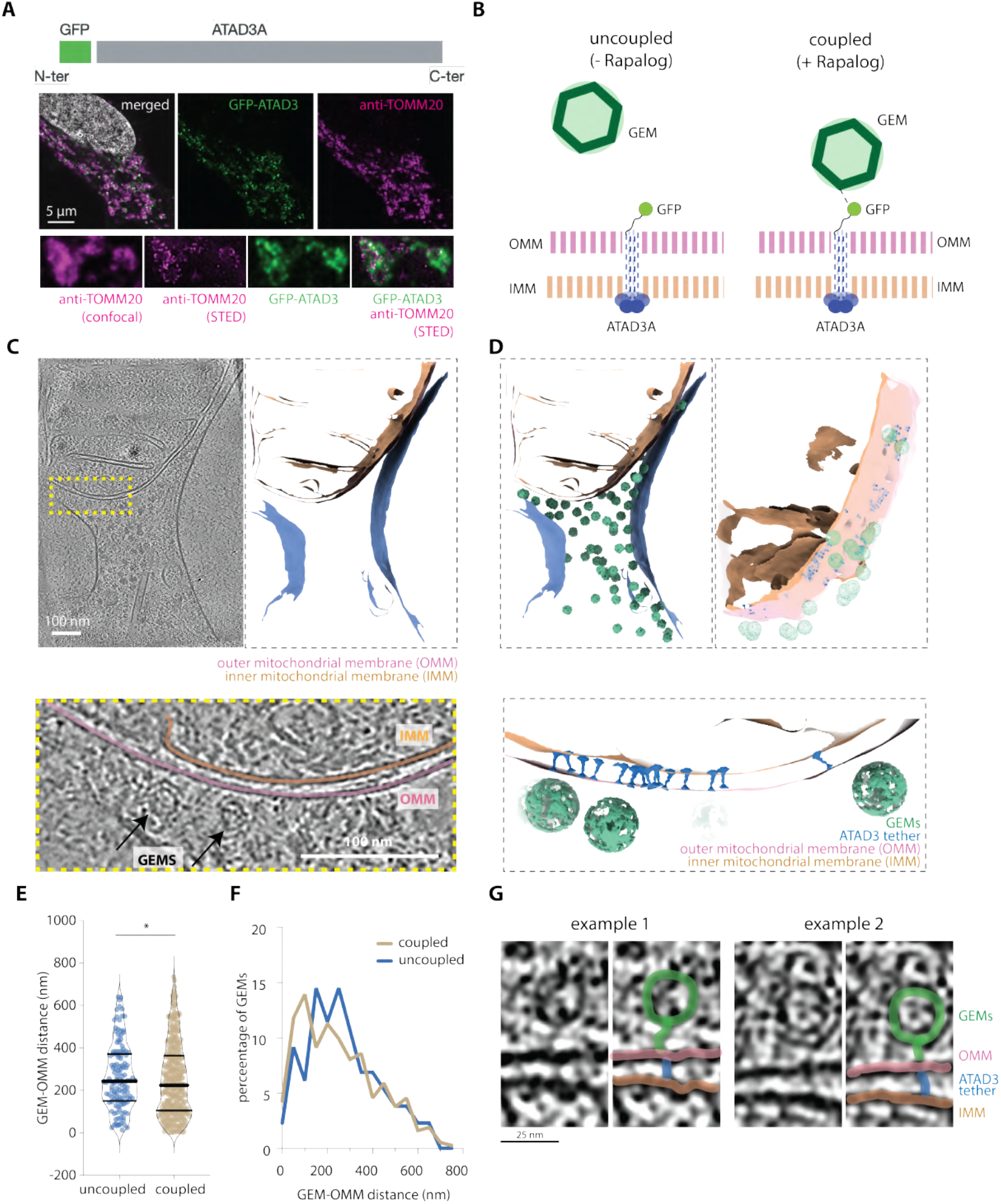
ATAD3A assembles into an intermembrane space-spanning tether. (**A**) Design of the N-terminus tagged GFP-ATAD3A construct, with representative confocal and STED images of cells expressing GFP-ATAD3A and stained with anti-TOMM20 showing colocalization of GFP with TOMM20. (**B**) Schematic representing the design of the GEM experiment. (**C**) A virtual slice from a 3D tomogram from a cell expressing GFP-ATAD3A and GEM donor plasmid treated with the coupler A/C heterodimerizer. Segmented membranes *(right)*. Zoomed-in view of inset (dashed yellow rectangle) with black arrows marking GEM particles next to OMM *(bottom)*. (**D**) Segmented membranes along with identified GEMs *(left)*. Zoomed in view showing identified GEM particles and ATAD3 tethers *(right and bottom)*. (**E**) The distances of identified GEMs from OMM, under both coupled and uncoupled conditions, are shown as a violin plot. Each data point represents an individual GEM particle. (**F**) Relative percentage of GEMs over GEM-OMM distance. (**G**) Two representative examples showing a GEM particle close to an intermembrane space-spanning tether with connecting density. Statistical significance in F is calculated using the KS test (131 GEMs in uncoupled, 373 GEMs in coupled) (\*\*\*\**p<* 0.0001, \*\*\**p<* 0.001, \*\**p<* 0.01, \**p<* 0.05).

In the resulting tomograms, we readily observe the characteristic ∼35 nm-diameter GEM throughout the cytoplasm (Fig. 4C, D) under both coupled and uncoupled conditions. We used an automated, pre-trained model to identify the location of each GEM particle with deep-learning-based segmentation software Fung et al. (2023); de Teresa-Trueba et al. (2023) (Fig. S5A) and performed subtomogram averaging to confirm that the detected particles formed the expected ∼35 nm icosahedral GEM (Fig. S5B–D). Using our surface morpho-metrics pipeline, we calculated the distance between each GEM particle and the OMM in cells co-expressing GFP-ATAD3A and the GEM tag in the presence (i.e., coupled) and absence (i.e., uncoupled) of rapalog treatment (Fig. 4D–F). We observed a significant reduction in the distance between the GEM particle and the OMM in the coupled condition relative to the uncoupled condition and an overall increase in the number of GEM particles positioned within 100 nm of the OMM in the coupled condition (Fig. 4E, F). Within these localized regions, we also observe GEM particles positioned next to clusters of tethers (Fig. 4D, G), with continuous connecting densities between the outer mitochondrial membrane and the GEM particles (Fig. 4G). Taken together, these results suggest that membrane-spanning tethers observed near replicating mtDNA are ATAD3A molecules positioned with their N-terminus exposed to the cytoplasm.

### ATAD3A remodels mitochondrial membranes

The results thus far point to a model in which ATAD3A forms intermembrane-spanning tethers that cluster specifically at sites of mtDNA replication. We sought to investigate whether this ATAD3A enrichment could promote the membrane remodeling we observed specifically at mtDNA replication sites by performing cryoET on cells overexpressing GFP-ATAD3A (ATAD3A *OE*) and ATAD3A *KD* cells (Fig. 5 and Fig. S6). In ATAD3A *OE* cells, we found that OMM-IMM distribution closely resembled that of mitochondria containing replicating mtDNA (Fig. 5A, B and Fig. 1H), with uniform distances across the IBM regions (Fig. 5C). Strikingly, in cells exhibiting very high levels of GFP-ATAD3A over-expression, we observed mitochondria completely devoid of cristae with multiple membranes stacked with uniform, ∼12 nm spacing (Fig. S6A, B). Within these stacked membranes, we also observe extensive intermembrane spanning densities resembling those identified by our GEM tags to be ATAD3A (Fig. S6C).

**Figure 5.**
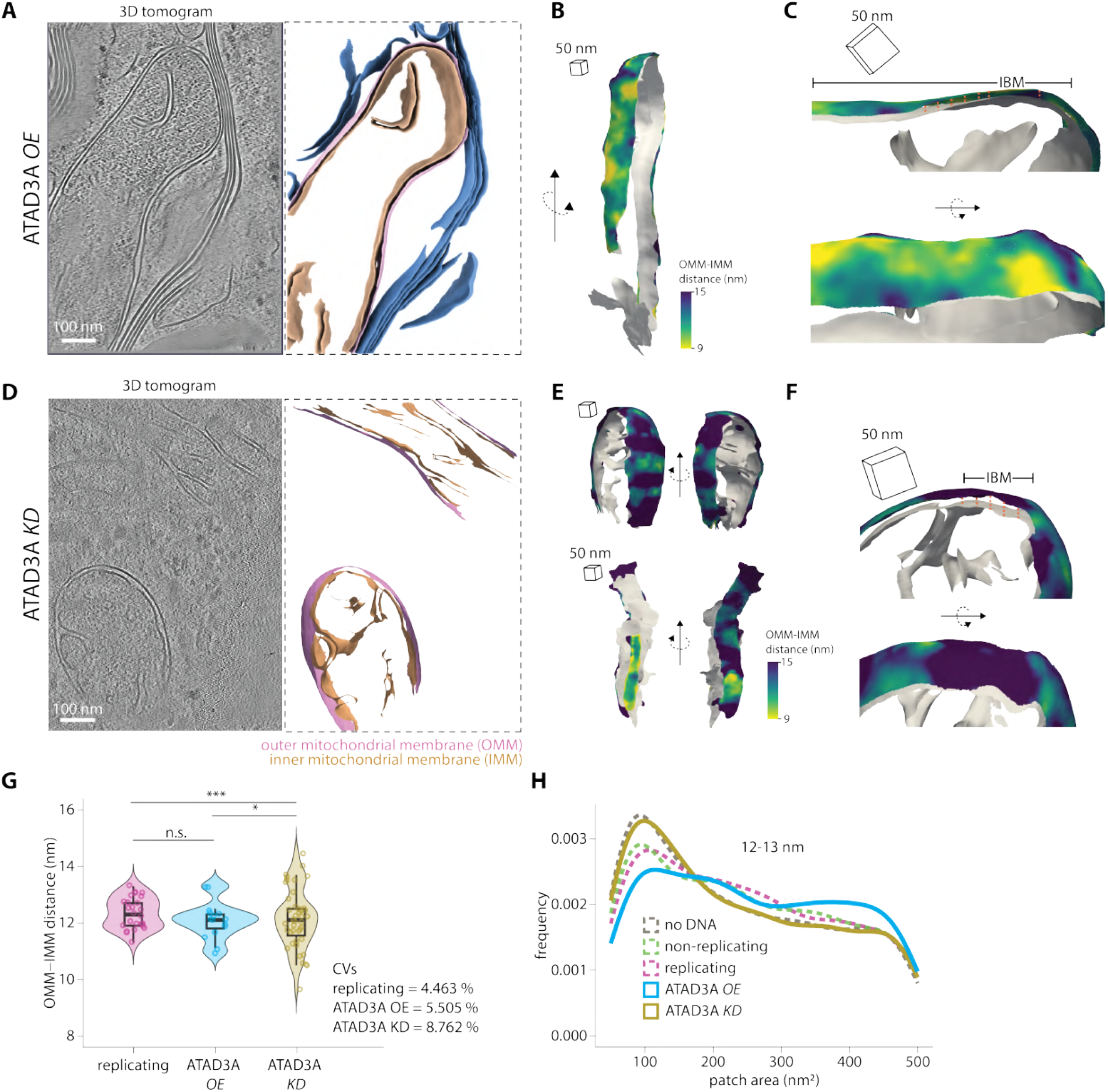
ATAD3A overexpression and knockdown remodels mitochondrial membranes. (**A**) Representative 3D tomogram from a cell overexpressing GFP-ATAD3A (ATAD3A *OE*). (**B**) Per-triangle OMM-IMM distance measurements mapped on the surface of the tomogram shown in A. (**C**) A zoomed-in view of mitochondria shown in B displaying consistent OMM-IMM distance in the IBM region. (**D**) Representative 3D tomogram of ATAD3A knockdown (ATAD3A *KD*) cells. (**E**) Per-triangle OMM-IMM distance measurements mapped on the surface for the tomogram shown in D. (**F**) A zoomed-in view of mitochondria shown in E displaying variable OMM-IMM distance in the IBM region. (**G**) Peak histogram values from each mitochondrion for OMM-IMM distance plotted as a violin plot for ATAD3A *OE* and ATAD3A *KD* cells, compared to mitochondria with replicating mtDNA (same as Fig. 1). Each dot represents the peak value for the corresponding mitochondrion. (*n* = 26 replicating, 15 = ATAD3A *OE*, 42 = ATAD3A *KD*). (**H**) Frequency distributions for the area of patches extracted after applying a threshold of 12–13 nm for the OMM-IMM distance. For statistical significance in G, coefficient of variation (CV) was calculated for each group and compared using the Feltz & Miller asymptotic test for equality of CVs. Pairwise comparisons were performed using a two-sample asymptotic Z-test, with Bonferroni correction applied for three comparisons (corrected *α* = 0.0167). (\*\*\*\**p<* 0.0001, \*\*\**p<* 0.001, \*\**p<* 0.01, \**p<* 0.05)

We observed the opposite effects of ATAD3A *KD*, with a significant increase in variability in OMM-IMM distances globally across the IMM and within the IBM regions(Fig. 5D–F). While we observe a tight distribution of OMM-IMM distances in ATAD3A *OE* cells, similar to “replicating” mtDNA containing mitochondria (Fig. 5G), this distribution is lost in ATAD3A *KD* cells, which displayed significantly higher variability in OMM-IMM distance (Fig. 5G). We extracted membrane patches containing 12-13 nm OMM-IMM distance (as in Fig. S1G, H) and observed a higher frequency of larger patches in ATAD3A *OE* cells and the opposite effect in ATAD3A *KD*, supporting the observation that OMM-IMM distance variability is lost under ATAD3A *OE* (Fig. 5H). Collectively, these results show that overexpression of ATAD3A promotes membrane remodeling toward fixed OMM-IMM spacing, which is lost in knockdown cells, suggesting that ATAD3A clustering at replicating mtDNA leads to membrane remodeling locally at these sites.

### ATAD3A is required for the recruitment of mitochondrial fission machinery at replicating mtDNA

On the basis of the results above, we propose that ATAD3A enrichment specifically at replicating mtDNA nucleoids promotes the recruitment of fission machinery and subsequent mitochondrial fission. A well-established marker of mitochondrial fission is the GT-Pase DRP1 Smirnova et al. (2001), which requires a critical threshold of dimer recruitment to the mitochondrial surface before assembling as a force-producing oligomeric ring to complete membrane scission. We observed that overexpression of ATAD3A led to an increase in DRP1 recruitment to mitochondria (Fig. 6A, B), whereas ATAD3A knockdown reduced the recruitment of DRP1 to mitochondria (Fig. 6C, D). These changes in DRP1 levels were also linked to changes in mitochondrial shape, with ATAD3A-overexpressing cells displaying fragmented networks and ATAD3A-knockdown cells displaying elongated mitochondria (Fig. 6E). We also observed more branched mitochondria in our cryo-ET datasets in ATAD3A knockdown cells (Fig. S7A). Although we observed a decrease in DRP1 recruitment in ATAD3A *KD* cells, we found that the amount of DRP1 associated with POLG2 relative to the entire mitochondrial network (Fig. 6F), does not significantly change under knockdown conditions (Fig. 6G, H). This would suggest that even under lower absolute levels of ATAD3A, the gradient across the mitochondrial network, with specific enrichment at replicating nucleoids, is maintained.

**Figure 6.**
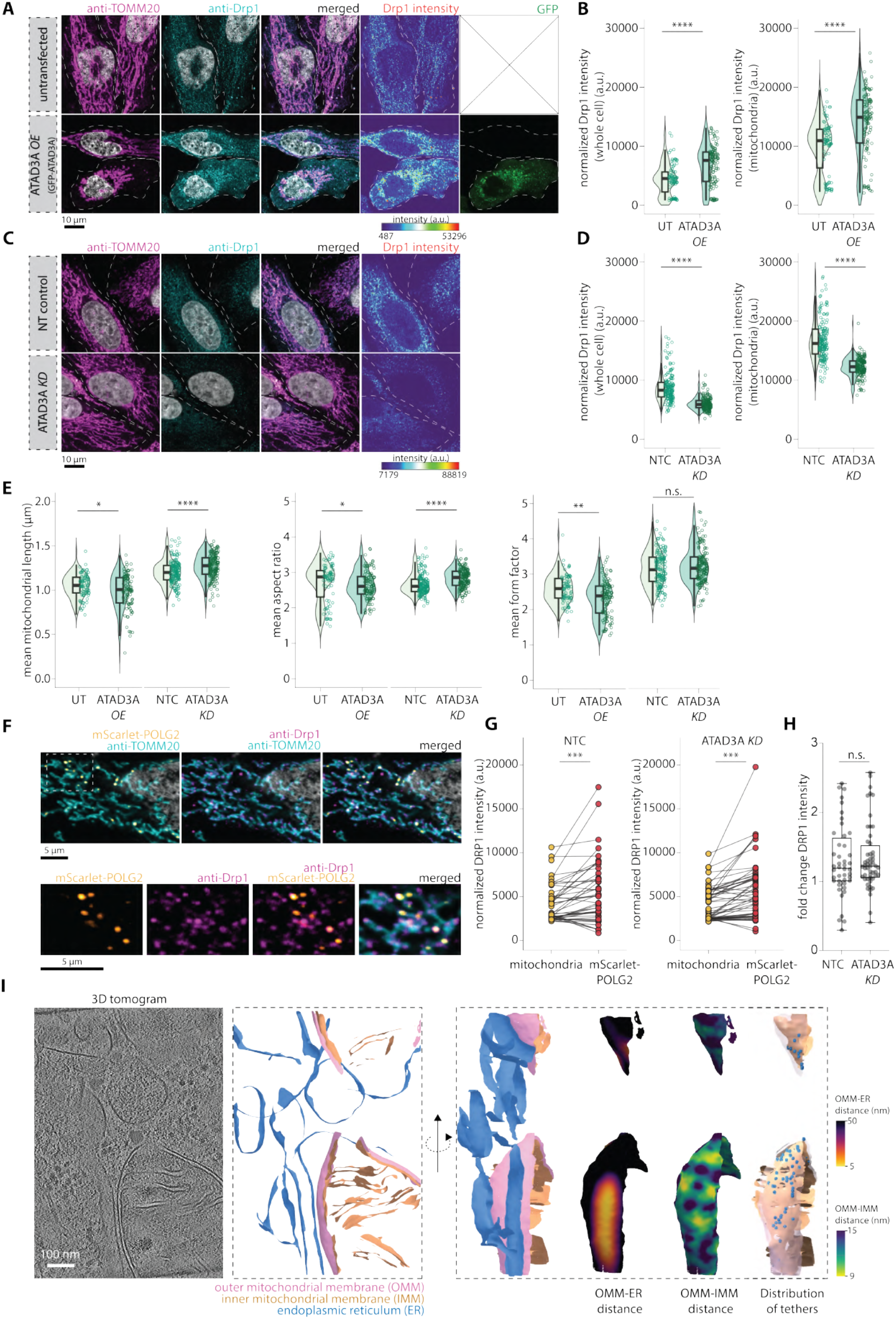
ATAD3A overexpression and knockdown remodels mitochondrial membranes. (**A**) Representative confocal images of HeLa cells ectopically expressing GFP-ATAD3A (ATAD3A *OE*) and stained with anti-TOMM20 and anti-DRP1 antibody. (**B**) Violin plots for whole cell DRP1 intensity normalized to cell area *(left)* or mitochondrial DRP1 intensity normalized to mitochondrial area *(right)*. Each dot represents one cell. (*n* = 75 cells in untransfected and 96 cells in ATAD3A *OE* from 3 biological replicates). (**C**) Representative confocal images of ATAD3A *KD* HeLa cells stained with anti-TOMM20 and anti-DRP1 antibody. (**D**) Violin plots for whole cell DRP1 intensity normalized to cell area *(left)* or mitochondrial DRP1 intensity normalized to mitochondrial area *(right)*. Each dot represents one cell. (*n* = 141 cells in NTC (no template control) and 134 cells in ATAD3A *KD* from 3 biological replicates). (**E**) Violin plots for mean mitochondrial length, mean aspect ratio and mean form factor for untransfected cells and ATAD3A *OE* cells (GFP-ATAD3A); and NTC (no template control) cells and ATAD3A *KD* cells. Each dot represents one cell. (*n* = 75 cells in untransfected and 86 cells in GFP-ATAD3A, *n* = 141 cells in NTC (no template control) and 134 cells in ATAD3A *KD* from 3 biological replicates). (**F**) Representative confocal images of ATAD3A *KD* HeLa cells transiently expressing mScarlet-POLG2, stained with anti-TOMM20 and anti-DRP1 antibody. (**G**) Pairwise plot showing normalized mitochondrial DRP1 intensity (normalized to mitochondrial area) and DRP1 intensity over mScarlet-POLG2 foci (normalized to mScarlet-POLG2 foci area) for both control (NTC) and ATAD3A *KD* cells. Each dot represents one cell (*n* = 44 cells in NTC (no template control) and 52 cells in ATAD3A *KD* from 2 biological replicates). (**H**) Fold change in DRP1 intensity (over mScarlet-POLG2 foci/mitochondria) in F is plotted as a box plot. Each dot represents one cell (*n* = 44 cells in NTC (no template control) and 52 cells in ATAD3A *KD* from 2 biological replicates). (**I**) Representative tomogram from the “replicating” mtDNA category showing close OMM-ER contact (*<* 15 nm), OMM-IMM distance, and distribution of ATAD3A tethers. Statistical significance in B, D, and G is calculated using the unpaired *t* -test. Statistical significance in F is calculated using a paired *t* -test. (\*\*\*\**p<* 0.0001, \*\*\**p<* 0.001, \*\**p<* 0.01, \**p<* 0.05)

Along with DRP1, binding of ER at sites of replication is important for fission Lewis et al. (2016); Friedman et al. (2011). Consistent with this, we found that a subpopulation of mitochondria with replicating mtDNA formed close contacts (*<* 15 nm) with ER (Fig. 6I and Fig. S7B, C), a well-established marker of mtDNA replication-linked fission. These close ER-OMM contacts formed in the same regions that had patches of uniform ∼12 nm OMM-IMM distance and were enriched with ATAD3A tethers (Fig. 6H and Fig. S7C). Taken together, these results suggest that ATAD3A is required for the recruitment of mitochondrial fission machinery at sites of mtDNA replication.

## Discussion

The signal linking mtDNA replication and mitochondrial fission, two processes separated by the barrier of two distinct mitochondrial membranes has remained unknown, despite their functional coupling being established nearly a decade ago Lewis et al. (2016). Using a correlative cryo-ET pipeline and quantitative analyses, we provide the first structural characterization of the local microenvironment surrounding mtDNA replication sites and identify the architectural features that underlie this coordination. Our findings reveal that replicating nucleoids are surrounded by distinct membrane ultrastructure, characterized by local remodeling driven by clustered, inter-membrane spanning tethers composed of ATAD3A. We show that this replication-dependent ATAD3A enrichment is both necessary and sufficient to recruit the mitochondrial fission machinery and induce fission. Together, these results establish ATAD3A as a structural scaffold that transmits information about the replication state of mtDNA across multiple mitochondrial sub-compartments to the cytoplasmic machinery that mediates fission.

While the interplay between large-scale mitochondrial membrane (especially cristae) remodeling and mtDNA has been established Kondadi and Reichert (2024), recent work has suggested that more localized changes can impact mtDNA signaling and function Newman et al. (2024); Zhang et al. (2026); Sen et al. (2022); He et al. (2022); Fu et al. (2023); Tábara et al. (2024). Such localized changes have been proposed in budding yeast cells, where each nucleoid exerts a local “sphere of influence” over surrounding membrane architecture Jakubke et al. (2021). We observe that mitochondria with replicating nucleoids contain uniform, ∼12 nm OMM-IMM distances relative to regions with non-replicating nucleoids or mtDNA-free regions (Fig. 1), demonstrating that membrane architecture is locally re-modelled in a replication-dependent manner. This is consistent with the “sphere of influence” model that proposes that mtDNA state (such as, replication) could dictate changes in the local microenvironment, which could serve as a structural cue for cytosolic machineries to sense the functional state of mtDNA nucleoids. Our cryo-ET data provides direct evidence confirming this model and identifies the specific membrane features, such as uniform OMM-IMM spacing, that define this zone.

A second defining feature of replicating nucleoid microenvironments is the enrichment of membrane-spanning tether densities in regions with consistent OMM-IMM spacing. Physical tethers between the IMM and OMM such as ATAD3A, have been proposed as mediators of communication between interior and exterior mitochondrial factors Ishihara et al. (2022); Zhao et al. (2019); Brar et al. (2024); Teng et al. (2016). Our study provides the first direct *in-situ* evidence that such tethers cluster specifically at sites of active mtDNA replication. Assigning molecular identities to specific densities within cellular cryo-ET data has long been a fundamental limitation of the field, but one that genetically encoded cryo-ET tags now make tractable. Using the GEM-tag technology Fung et al. (2023), we identify these tethers as ATAD3A, the AAA+ ATPase whose N-terminal domain faces the cytoplasm and whose c-terminal ATPase domain is matrix localized. This unique topology positions ATAD3A to span the intermembrane space and interact with both mitochondrial matrix and cytoplasmic components. Consistent with this, ATAD3A had separately been shown to interact with both mtDNA and DRP1 Zhao et al. (2019); He et al. (2007). We propose that replication-associated decompaction of mtDNA Brüser et al. (2021) exposes nucleoid binding sites that drive local ATAD3A clustering, either through direct DNA binding or via nucleoid-associated proteins such as TFAM Ishihara et al. (2022) (Fig. S7D). The local accumulation of ATAD3A would then result in the local “zippering” of IMM to OMM across the intermembrane space at the mtDNA replication site, producing the consistent inter-membrane distances we observed in our cryo-ET data.

The functional consequence of local membrane remodeling and ATAD3A enrichment at mtDNA replication sites is the localized recruitment of mitochondrial fission machinery. We show that ATAD3A-enriched regions display increased association with ER membranes and DRP1, two well-established markers of mitochondrial fission. We show that ATAD3A overexpression enhances this association, whereas knockdown reduces this recruitment, directly linking ATAD3A levels to fission factor recruitment and completion at the local level. These findings provide a structural basis for prior observations that ATAD3A knockdown simultaneously impairs fission and disrupts nucleoid distribution, and that ATAD3A is required for replication-stress-induced mtDNA release, unifying these phenotypes under a single mechanism in which ATAD3A physically couples replication state to membrane remodeling.

The concept of ATAD3A-dependent fission recruitment is reminiscent of how bacteria position their division septum, where MinCDE gradients restrict Z-ring assembly by locally concentrating division factors above a threshold required for productive septation Wu and Errington (2012). We propose that ATAD3A performs an analogous function in mitochondria, wherein its mtDNA replication-dependent enrichment at adjacent membranes creates a local concentration of N-terminal DRP1-binding sites that exceeds the threshold required for productive fission (Fig. S7D). This is consistent with the observation that not all DRP1 recruitment events lead to membrane scission, and that DRP1 must accumulate above a critical concentration to drive fission Ji et al. (2015); Zollo et al. (2026). In this model, the ATAD3A gradient generated by mtDNA replication state serves as a structural scaffold, ensuring that the fission threshold is preferentially enriched at replication sites rather than at random points along the mitochondrial network, thereby coupling the timing and position of fission to mtDNA replication.

Taken together, our results support a model in which mtDNA replication-dependent accumulation of ATAD3A at adjacent membranes creates nanoscale IMM-OMM tethers that remodel the surrounding microenvironment and position the fission machinery at the replication site for downstream fission. This mechanism resolves a long-standing question about how two processes that occur in distinct organellar compartments are spatially coordinated. More broadly, these findings establish a structural paradigm for inter-compartment organellar communication, wherein the functional state of a process on one side of a membrane is encoded in the nanoscale architecture of the bounding membrane and read out as a recruitment platform on the other.

## Limitations of study

While this study establishes the structural basis for ATAD3A-dependent coupling of mtDNA replication and fission, several important questions remain. All experiments in this study were performed in HeLa cells; therefore, determining whether this mechanism operates similarly in post-mitotic cell types, such as neurons, where ATAD3A variants cause neurodegenerative disease, will be an important avenue for future investigation. Additionally, our experiments rely on the transient expression of mScarlet-POLG2, which inevitably leads to higher POLG2 levels than endogenous levels. Whether mtDNA replication stress engages a similar structural pathway altering ATAD3A tether assembly,local membrane architecture, and fission coupling remains unknown. Given that ATAD3A is required for replication stress-induced mtDNA release, understanding how replication stress modifies this structural scaffold may reveal how the replication-fission axis contributes to organellar quality control.

## Resource Availability

### Lead Contact

Danielle A. Grotjahn

### Material availability

Any requests for materials from this study should be addressed to Danielle A. Grotjahn.

### Data and code availability

Maps have been deposited in EMDB under accession code EMD-77685. The tilt-series micrograph data and other associated cryo-ET files have been deposited in EMPIAR under accession code EMPIAR-13702. This paper does not report original code.

## Acknowledgements

We thank William Lessin at The Scripps Research Institute Hazen cryo-electron microscopy facility for microscope support. We thank Jean-Christophe Ducom and Lisa Dong at The Scripps Research Institute for computational support. We thank Brian Seegers and Alphonse Owirka at The Scripps Research Institute Flow Cytometry facility for FACS sorting support. We thank Herman Fung for his input on GEM experiment design and analysis. We also thank the members of the Grotjahn Lab for their critical input on the manuscript. D.P is supported Scripps Research start-up funds. D.A.G. is supported by The Pew Scholars Program, Nadia’s Gift Foundation Innovator Award of the Damon Runyon Cancer Foundation (DRR-65-21), and the National Institutes of Health (NIH) grant (R21NS142779). R.LW. is supported by the National Institutes of Health (NIH) grant RF1NS125674. This work used equipment supported by NIH grant S10OD032467. Assistance from ChatGPT-5.2 (Open AI, https://chat.openai.com/), Claude (Anthropic), and Grammarly’s AI tool was utilized to improve the clarity, grammar, and conciseness of the manuscript text.

## Author Contributions

**N. Dua:** Conceptualization, Data curation, Formal analysis, Investigation, Methodology, Software, Validation, Visualization, Writing - original draft, Writing - reviewing and editing.

**B. Ma:** Formal analysis, Writing – reviewing, and editing.

**S. Oviedo:** Formal analysis, Investigation, Writing – reviewing, and editing.

**H. Rahmani:** Data curation, Software, Validation, Writing - reviewing and editing.

**T. Boyd:** Formal analysis, Writing reviewing and editing.

**D. Park:** Funding acquisition, Supervision, Writing - reviewing and editing.

**R.L Wiseman:** Funding acquisition, Supervision, Writing - reviewing and editing.

**D. A. Grotjahn:** Conceptualization, Funding acquisition, Methodology, Project administration, Resources, Supervision, Visualization, Writing - original draft, Writing - reviewing and editing.

## Methods

### Cell lines and growth conditions

HeLa cells and U2OS cells were grown in Dulbecco’s Modified Eagle Medium (DMEM) + GlutaMax supplemented with 10% FBS and Glutamine (4 mM). Cells were incubated at 37°C with 5% CO_2_ and utilized for experiments only up to passage 23 and were regularly tested for mycoplasma. For transfections, cells were grown to ∼70% confluency and transfected with Lipofectamine 3000 (Thermo Fisher Scientific).

### Grid preparation

#### SYBR Gold and mScarlet-POLG2 experiments

HeLa cells were transfected with a plasmid expressing mScarlet-POLG2 at ∼70% confluency in a 10 cm dish. The transfection media was replaced with fresh media after 4 hours and the cells were allowed to incubate overnight. The following day, cells were trypsinized with 1 ml trypsin and resuspended in 4 ml of sorting buffer (1X PBS Ca/Mg^++^ free), 2.5 mM EDTA, 25 mM HEPES (pH 7.0), 1% FBS (Heat inactivated), 1% Pen-Strep). The cell suspension was kept on ice until sorting into 6 well plates containing EM grids. Quantifoil R1/4 or 2/2 Holey Carbon Grids were UV sterilized, and plasma cleaned using Solarus plasma cleaner for 7 seconds. Grids were then coated with fibronectin (0.5 mg/ml) by placing them inverted on a droplet containing a 1:1 mix of fibronectin (1 mg/ml) and PBS for 15 minutes. The grids were then transferred to PBS and then immediately placed in a 6 well plate and 1 ml of fresh culture media was added to each well. 50,000 to 70,000 cells were sorted into each well containing grids, gated to select for cells positive for mScarlet fluorescence. Post sorting, the plates were incubated for ∼8 hours. Prior to vitrification, the cells were stained with SYBR Gold. For this, 0.5 *µ*l/ml of SYBR Gold (10,000X stock in DMSO, Thermo Fisher Scientific) was prepared in fresh culture media and prewarmed. Culture media from 6 well plate was aspirated and replaced with media containing SYBR Gold and incubated for 6–8 minutes. Cells were then washed twice with prewarmed culture media and the grids were taken for vitrification.

#### GEM experiments

Quantifoil R1/4 or 2/2 Holey Carbon Grids were used for cryo-ET experiments. The grids were UV sterilized, and plasma cleaned using Solarus plasma cleaner for 7 seconds. Grids were then coated with fibronectin (0.5 mg/ml) by placing them inverted on a droplet containing a 1:1 mix of fibronectin (1 mg/ml) and PBS for 15 minutes. The grids were then transferred to PBS and then immediately placed in custom grid holders inside a MatTek dish. HeLa cells were trypsinized from a 10 cm plate by aspirating used media, washing with 2 ml PBS and then adding 1 ml Trypsin-EDTA. Cells were incubated with Trypsin for ∼5 mins, then resuspended in 9 ml fresh media. In a separate tube, ∼ 35 *µ*l of this cell suspension was mixed with 265 *µ*l of fresh media, the final 300 *µ*l of cell suspension was then placed on top of the grids placed in the grid holder in MatTek dishes. Cells were allowed to adhere to the grids for 30 min, and then 2 ml of fresh media was added to the MatTek dishes and incubated. Grids were incubated for ∼ 8 hours and then cells were transfected with GFP-ATAD3 and GEM donor plasmid. The transfection mix was removed after 4 hours and 2 *µ*g/ml doxycycline was added after 16 hours of overnight incubation. The cells were incubated for 24 hours and then treated with 0.5 *µ*M A/C heterodimerizer (Takara) and 50 nM Halo tag (HT105A) ligand for 1 hour before vitrification.

For vitrification, grids were plunge frozen in an ethane/propane mixture using a Vitrobot Mark 4 (Thermo Fisher Scientific). The Vitrobot chamber was maintained at 37°C and 90% relative humidity. Blotting was performed manually using a Whatman #1 filter paper from the side port of the Vitrobot chamber for 12–15 seconds. Post-vitrification, grids were clipped in Cryo-FIB Autogrid (Thermo Fisher Scientific).

### Cryo-fluorescence microscopy

Pre-Cryo-FIB milling fluorescence images were acquired on a Leica CryoCLEM widefield microscope. A montage of fluorescence and brightfield image z-stacks (∼25 *µ*m) was acquired using the Matrix Maps feature of Leica LasX acquisition software. The microscope is equipped with a sola light engine light source, and filter cubes for GFP (470/40 Dichroic, 495 Emission 525/50), RFP (546/10 Dichroic, LP 560 emission BP 585/40) and Y5 (620/60 excitation, 660 emission 700/75) and a Leica DFC9000 GT camera with HC PL APO 50 ×/NA 0.9 objective. Post-cryo-FIB images were acquired with a METEOR in-chamber fluorescence imaging system driven by Odemis version 3.6.3 software Smeets et al. (2021). Z-stacks of thickness 4 *µ*m with step size 0.5 *µ*m were acquired at each milled lamella site. Correlation between EM and FM images was performed using eC-CLEM plugin in Icy Software Heiligenstein et al. (2017). Briefly, the fluorescence image acquired using METEOR was correlated to the SEM image of the lamella. Simultaneously, high magnification polygon montages were correlated to the low magnification TEM images of the lamella. The transformed fluorescence and SEM images were also correlated to the low magnification TEM images. Only rigid body transformations were allowed for all the correlations.

### Cryo-FIB milling

Cryo-FIB milling was performed on an Aquilos dual-beam cryo-FIB/SEM microscope (Thermo Fisher Scientific). Prior to milling, grids were coated with an organoplatinum layer using a gas injection system for 30 seconds. A sputter coat was applied for 15 seconds (1 kV, 20 mA, 10 Pa). Milling was performed using AutoTEM as previously described Barad et al. (2023). After the automatic rough milling and thinning were completed, lamella were polished to a final thickness of ∼150– 200 nm using an ion beam of 50 pA. Upon completion of polishing, a final layer of sputter coating was applied for 22 seconds (1 kV, 10 mA, 10 Pa).

### Tilt series data collection

Tilt series acquisition was performed on a 300 keV Titan Krios microscope (Thermo Fisher Scientific), equipped with a K3 Summit direct electron camera and a BioQuantum energy filter (Gatan). SerialEM Mastronarde (2005) was used to acquire a low dose (1 e^−^/Å^2^) TEM image at 33k magnification at each lamella site. These images allow us to visualize cellular features, including mitochondria, with characteristic inner and outer membranes. Multiple tilt series were acquired using a Python based parallel cryo-ET (PACE-tomo) Eisenstein et al. (2023) data collection at a magnification of 53k and a pixel size of 1.662 Å and a defocus range of −4 to −6. The grid was tilted −60°to +60°in 3°steps centered on a pre-tilt of −10°, and the tilts were acquired with dose fractionation at 0.3 e^−^/Å^2^ per frame, with 10 frames each tilt leading to a total dose of 3 e^−^/Å^2^ per tilt.

### Tilt series reconstruction and processing

CTF estimation and motion correction on dose-fractionated tilt series was performed in Warp, and an averaged tilt series was generated for alignment Tegunov and Cramer (2019). For tilt series alignment, either a patch tracking model or a bead tracking model (using fiducials from sputter coating) was used in etomo Mastronarde and Held (2017). The resulting alignment files were imported into Warp to generate tomostar files, which were dose-adjusted using adjust_tomostar.py (https://github.com/GrotjahnLab/warp_lamella_adapters). Final tomograms were reconstructed in Warp, at bin 6 (pixel size 9.98 Å) or bin 8 (pixel size 13.30 Å). During reconstruction, half maps were generated for denoising in Deepdewedge Wiedemann and Heckel (2024). A Deep-dewedge model was trained on 2 selected tomograms with a subtomogram size of 72 pixels. The trained model was then used to refine all the tomograms in the dataset.

### Membrane segmentations and Surface morphometrics

Membrane segmentations were performed on denoised tomograms at a pixel size of 9.98 Å using MembrainSeg Lamm et al. (2025). The segmentations were then imported into AMIRA for manual correction and annotation of distinct membranes, such as OMM, IMM and ER. These annotations were exported as different label files for further morphometrics analysis. For each label file, segmentations were first converted to membrane meshes followed by pycurv analysis, as described previously Barad et al. (2023).

### GEMs particle picking and subtomogram averaging

We used a pretrained model Fung et al. (2023) in Deepict de Teresa-Trueba et al. (2023) to locate GEM particles in our tomograms. We ran DeePiCT predictions on both, warp-generated tomograms at a pixel size of 13.3 Å, and the same tomograms filtered in eman2 Tang et al. (2007) with the following parameters: `filter.low-pass.gauss:cutoff_abs=0.25,filter.highpass.gauss:cutoff_pixels=5, normalize,threshold.clampminmax.nsigma: nsigma=3`. The predictions were filtered with a 0.5 threshold, and then the two sets of particles were combined and duplicates removed using a python code. The combination particle sets were then imported in Cube and obvious false-positives (particles inside mitochondria or other membrane compartments, outside of the tomogram, and the ones located on frost) were removed manually. The rest of these particles were then assigned random Euler angles and imported into i3. We performed 1 cycle of translational only and 3 cycles of translational and rotational alignments in i3. All the remaining particles were retained for final alignment since no obvious false positives were identified upon classification (Fig. S5C). The alignment information (in the trf format) was then extracted and converted into star files for further in-context analysis.

### Clustering analysis

To assess clustering of ATAD3 tethers, we utilized Tomospatstat Martin-Solana et al. (2024), which employs Ripley’s K function *K*(*r*) to determine the number of tethers present within a given radius *r*. We plot the ratio *K*(*r*)*/K*_csr_(*r*), where *K*_csr_(*r*) represents a value expected from complete spatial randomness (csr). In a scenario where there is no clustering, this ratio would be equal to 1. We combine the maximum value of *K*(*r*)*/K*_csr_(*r*) for different radii intervals from multiple tomograms and plot them as violin plots. The identified coordinates for tethers resulting from blinded manual particle picking were used as input for this analysis.

### Tether vs random patch analysis

Tether membrane patches were identified by using the starfile containing coordinates for manually identified tethers. We used find_IMM_patches_for_ATP_synthase.py script (https://github.com/GrotjahnLab/patch_analysis/blob/main/find_IMM_patches_for_ATP_synthase.py) to find the coordinates and identifiers of the nearest OMM surface triangle and generate random patches. Subsequently, the corresponding tether membrane patch was defined by all the triangles within a 10 nm distance.

### ATAD3A tomogram simulation

The AlphaFold model of the ATAD3A and PDB files of OMM and IMM from CHARMM were assembled in ChimeraX Pettersen et al. (2021) and all the water molecules were removed. We expanded each membrane 5 × 5 and combined the PDB files. We generated an MRC model at pixel size of 3.32 Å of ATAD3A model between the membranes using MolMap function in ChimeraX. The volume generated using molmap has values between 0 and 1. To match the contrast with simulated tomograms we multiplied every voxel value by 1000 and then added random gaussian noise with a mean of 100 and standard deviation of 25. The resulting MRC model was binned by a factor of 3, to be imported in Cryotomosim Purnell et al. (2023) to create a simulated tomogram. For simulation, we generated a 400×400 ×100 tomogram, using tilts of *±*60° with a step size of 3° and defocus of − 4 *µ*m. Dose damage and tilt error were turned off to increase contrast.

### Fluorescence microscopy and image analysis

Cells were fixed with prewarmed 4% Image-IT paraformaldehyde fixative solution (Thermo Fisher Scientific) for 20 minutes at room temperature (RT). Fixation was quenched with a 0.1M Glycine solution prepared in PBS for 5 minutes with gentle shaking. Cells were then permeabilized using 0.1% Triton-X 100 in PBS for 10 minutes. After permeabilization, blocking solution (0.1% Tween-20 and 1% BSA in PBS) was added to the coverslips and incubated at RT for 1 hour with gentle shaking. This was followed by incubation with primary antibody prepared in blocking solution for 1 hr at RT with gentle shaking. Subsequently, coverslips were washed thrice with PBS (each wash for 8 min) and probed with the secondary antibody prepared in blocking solution. The coverslips were washed again with PBS and mounted using SlowFade Diamond Antifade soft-setting mountant with DAPI. Antibodies used: anti-ATAD3A (Sigma-Aldrich; HPA065305; 1:250 dilution), anti-DNA (Sigma-Aldrich; CBL186 anti DNA antibody clone AC-30-10; 1:250 dilution), anti-TOMM20 (AbCam; ab56783; 1:500 dilution); anti-Drp1 (Proteintech; 129571-1-AP; 1:250 dilution), Star-Red secondary antibody (Abberior, NC1933870, 1:1000 dilution), Star-Orange secondary antibody (Abberior, NC1933866, 1:1000 dilution), Alexa Fluor 488 anti-Mouse IgG secondary antibody (AbCam; ab150113; 1:1000 dilution).

Confocal imaging was performed on a Zeiss LSM 880 confocal microscope equipped with a 32-channel array detector (Quasar), 2 side PMT detectors and a 63 ×Plan-Apochromat NA 1.4 objective. Images were acquired at a pixel size of 0.07 *µ*m. For STED imaging, the Abberior Facility Line 3D STED “super-resolution” microscope equipped with two pulsed STED depletion lasers (775 nm & 595 nm) and spectral detection (3 APD detectors and 1 Matrix detector) and an Olympus 60 × UPLanXApo NA 1.42 objective was used. Images were acquired at a pixel size of 0.02 *µ*m.

Images were deconvolved using Huygens Professional software for visualization and segmentation. All image analysis was performed in Fiji. For the segmentation of mitochondria, a custom pipeline was set up in Fiji, briefly involving the following steps: (a) background subtraction (rolling ball radius: 50 pixels); (b) sigma filter plus (c) enhanced local contrast (CLAHE); (d) tubeness filter (e) images conversion to 8-bit; (f) thresholding (g) images conversion to mask, despeckled, and outlier removal. Mitochondrial descriptors (branch length, area, and number of junctions) were obtained using MitoAnalyzer Chaudhry et al. (2020). For detecting clusters in STED resolved images, ‘Find Maxima’ function with a prominence score of *>* 5 was used, and then the SSIDC cluster indicator in BioVoxxel Toolbox was used with a distance of 0.5 *µ*m and minDensity of either 5 or 15 particles Brocher (2026).

## References

Abramson, J. et al. (2024). Accurate structure prediction of biomolecular interactions with AlphaFold 3. Nature, 630:493–500.

Arguello, T. et al. (2021). ATAD3A has a scaffolding role regulating mitochondria inner membrane structure and protein assembly. Cell Reports, 37:110139.

Ban-Ishihara, R., Ishihara, T., Sasaki, N., Mihara, K., and Ishihara, N. (2013). Dynamics of nucleoid structure regulated by mitochondrial fission contributes to cristae reformation and release of cytochrome c. Proceedings of the National Academy of Sciences, 110: 11863–11868.

Barad, B. A., Medina, M., Fuentes, D., Wiseman, R. L., and Grotjahn, D. A. (2023). Quantifying organellar ultrastructure in cryo-electron tomography using a surface morphometrics pipeline. Journal of Cell Biology, 222:e202204093.

Brar, K. K., Hughes, D. T., Morris, J. L., Subramanian, K., Krishna, S., Gao, F., Rieder, L. S., Uhrig, S., Freeman, J., Smith, H. L., Jukes-Jones, R., Avezov, E., Nunnari, J., Prudent, J., Butcher, A. J., and Mallucci, G. R. (2024). PERK-ATAD3A interaction provides a subcellular safe haven for protein synthesis during ER stress. Science, 385(6712):eadp7114.

Brocher, J. biovoxxel/BioVoxxel-Toolbox, (2026). URL https://github.com/biovoxxel/BioVoxxel-Toolbox.

Brüser, C., Keller-Findeisen, J., and Jakobs, S. (2021). The TFAM-to-mtDNA ratio defines inner-cellular nucleoid populations with distinct activity levels. Cell Reports, 37:110000.

Chaudhry, A., Shi, R., and Luciani, D. S. (2020). A pipeline for multidimensional confocal analysis of mitochondrial morphology, function, and dynamics in pancreatic ω-cells. American Journal of Physiology-Endocrinology and Metabolism, 318:E87–E101.

Cooper, H. M., Yang, Y., Ylikallio, E., Khairullin, R., Woldegebriel, R., Lin, K. L., Euro, L., Palin, E., Wolf, A., Trokovic, R., Isohanni, P., Kaakkola, S., Auranen, M., Lönnqvist, T., Wanrooij, S., and Tyynismaa, H. (2017). ATPase-deficient mitochondrial inner membrane protein ATAD3A disturbs mitochondrial dynamics in dominant hereditary spastic paraplegia. Human Molecular Genetics, 26(8):1432–1443.

Craven, L., Alston, C. L., Taylor, R. W., and Turnbull, D. M. (2017). Recent advances in mitochondrial disease. Annu. Rev. Genom. Hum. Genet., 18:257–275.

de Teresa-Trueba, I. et al. (2023). Convolutional networks for supervised mining of molecular patterns within cellular context. Nat Methods, 20:284–294.

Eisenstein, F. et al. (2023). Parallel cryo electron tomography on in situ lamellae. Nat Methods, 20:131–138.

Filograna, R., Mennuni, M., Alsina, D., and Larsson, N.-G. (2021). Mitochondrial dna copy number in human disease: the more the better? FEBS Letters, 595(8):976–1002. doi: 10.1002/1873-3468.14021.

Friedman, J. R. et al. (2011). Er tubules mark sites of mitochondrial division. Science, 334: 358–362.

Fu, Y., Sacco, O., DeBitetto, E., Kanshin, E., Ueberheide, B., and Sfeir, A. (2023). Mitochondrial DNA breaks activate an integrated stress response to reestablish homeostasis. Molecular Cell, 83(20):3740–3753.e9.

Fung, H. K. H. et al. (2023). Genetically encoded multimeric tags for subcellular protein localization in cryo-EM. Nat Methods, 20:1900–1908.

Garrido, N. et al. (2003). Composition and dynamics of human mitochondrial nucleoids. MBoC, 14:1583–1596.

Gilquin, B. et al. (2010). The AAA+ ATPase ATAD3A controls mitochondrial dynamics at the interface of the inner and outer membranes. Molecular and Cellular Biology, 30: 1984–1996.

Harel, T., Yoon, W. H., Garone, C., Gu, S., Coban-Akdemir, Z., Eldomery, M. K., Posey, J. E., Jhangiani, S. N., Rosenfeld, J. A., Cho, M. T., Fox, S., Withers, M., Brooks, S. M., Chiang, T., Duraine, L., Erdin, S., Yuan, B., Shao, Y., Moussallem, E., Lamperti, C., others, and Lupski, J. R. (2016). Recurrent de novo and biallelic variation of ATAD3A, encoding a mitochondrial membrane protein, results in distinct neurological syndromes. American Journal of Human Genetics, 99(4):831–845.

He, B. et al. (2022). Mitochondrial cristae architecture protects against mtDNA release and inflammation. Cell Reports, 41:111774.

He, J. et al. (2007). The AAA+ protein ATAD3 has displacement loop binding properties and is involved in mitochondrial nucleoid organization. Journal of Cell Biology, 176:141–146.

Heiligenstein, X., Paul-Gilloteaux, P., Raposo, G., and Salamero, J. (2017). ec-CLEM: A multidimension, multimodel software to correlate intermodal images with a focus on light and electron microscopy. Methods Cell Biol, 140:335–352.

Ishihara, T., Ban-Ishihara, R., Ota, A., and Ishihara, N. (2022). Mitochondrial nucleoid trafficking regulated by the inner-membrane aaa-atpase atad3a modulates respiratory complex formation. Proceedings of the National Academy of Sciences, 119(e2210730119).

Jakubke, C. et al. (2021). Cristae-dependent quality control of the mitochondrial genome. Science Advances, 7:eabi8886.

Ji, W. K., Hatch, A. L., Merrill, R. A., Strack, S., and Higgs, H. N. (2015). Actin filaments target the oligomeric maturation of the dynamin GTPase Drp1 to mitochondrial fission sites. eLife, 4:e11553.

Jo, S., Kim, T., Iyer, V. G., and Im, W. (2008). CHARMM-GUI: a web-based graphical user interface for CHARMM. Journal of Computational Chemistry, 29(11):1859–1865.

Kleele, T. et al. (2021). Distinct fission signatures predict mitochondrial degradation or biogenesis. Nature, 593:435–439.

Kondadi, A. K. and Reichert, A. S. (2024). Mitochondrial dynamics at different levels: From cristae dynamics to interorganellar cross talk. Annu. Rev. Biophys., 53:annurev–biophys–030822–020736.

Lamm, L. et al. (2025). MemBrain v2: an end-to-end tool for the analysis of membranes in cryo-electron tomography. Preprint at bioRxiv. doi: 10.1101/2024.01.05.574336.

Lewis, S. C., Uchiyama, L. F., and Nunnari, J. (2016). Er-mitochondria contacts couple mtdna synthesis with mitochondrial division in human cells. Science, 353.

Liu, T., Stephan, T., Chen, P., Keller-Findeisen, J., Chen, J., Riedel, D., Yang, Z., Jakobs, S., and Chen, Z. (2022). Multi-color live-cell sted nanoscopy of mitochondria with a gentle inner membrane stain. Proceedings of the National Academy of Sciences, 119 (52):e2215799119. doi: 10.1073/pnas.2215799119.

Martin-Solana, E. et al. (2024). Progressive alterations in polysomal architecture and activation of ribosome stalling relief factors in a mouse model of Huntington’s disease. Neurobiol Dis, 195:106488.

Mastronarde, D. N. (2005). Automated electron microscope tomography using robust prediction of specimen movements. Journal of Structural Biology, 152:36–51.

Mastronarde, D. N. and Held, S. R. (2017). Automated tilt series alignment and tomographic reconstruction in IMOD. J Struct Biol, 197:102–113.

Miyakawa, I. (2017). Organization and dynamics of yeast mitochondrial nucleoids. Proc Jpn Acad Ser B Phys Biol Sci, 93:339–359.

Newman, L. E. et al. (2024). Mitochondrial dna replication stress triggers a pro-inflammatory endosomal pathway of nucleoid disposal. Nat Cell Biol, 26:194–206.

Peralta, S., Goffart, S., Williams, S. L., Diaz, F., Garcia, S., Nissanka, N., Area-Gomez, E., Pohjoismäki, J., and Moraes, C. T. (2018). ATAD3 controls mitochondrial cristae structure in mouse muscle, influencing mtDNA replication and cholesterol levels. Journal of Cell Science, 131(13):jcs217075.

Pettersen, E. F., Goddard, T. D., Huang, C. C., Meng, E. C., Couch, G. S., Croll, T. I., Morris, J. H., and Ferrin, T. E. (2021). UCSF ChimeraX: Structure visualization for researchers, educators, and developers. Protein Science, 30(1):70–82.

Puchades, C., Sandate, C. R., and Lander, G. C. (2020). The molecular principles governing the activity and functional diversity of AAA+ proteins. Nature Reviews Molecular Cell Biology, 21(1):43–58.

Purnell, C., Heebner, J., Swulius, M. T., Hylton, R., Kabonick, S., Grillo, M., Grigoryev, S., Heberle, F., Waxham, M. N., and Swulius, M. T. (2023). Rapid synthesis of cryo-ET data for training deep learning models. bioRxiv, page 2023.04.28.538636.

Sen, A. et al. (2022). Mitochondrial membrane proteins and VPS35 orchestrate selective removal of mtDNA. Nat Commun, 13:6704.

Smeets, M., Bieber, A., Capitanio, C., Schioetz, O., van der Heijden, T., Effting, A., Piel, É., Lazem, B., Erdmann, P., and Plitzko, J. (2021). Integrated cryo-correlative microscopy for targeted structural investigation in situ. Microscopy Today, 29(6):20–25.

Smirnova, E., Griparic, L., Shurland, D. L., and van der Bliek, A. M. (2001). Dynamin-related protein Drp1 is required for mitochondrial division in mammalian cells. Molecular Biology of the Cell, 12(8):2245–2256.

Stephan, T., Roesch, A., Riedel, D., and Jakobs, S. (2019). Live-cell STED nanoscopy of mitochondrial cristae. Scientific Reports, 9(1):12419. doi: 10.1038/s41598-019-48838-2.

Tábara, L. C. et al. (2024). MTFP1 controls mitochondrial fusion to regulate inner membrane quality control and maintain mtDNA levels. Cell, 187:3619–3637.e27.

Tang, G. et al. (2007). EMAN2: an extensible image processing suite for electron microscopy. J Struct Biol, 157:38–46.

Taylor, R. W. and Turnbull, D. M. (2005). Mitochondrial dna mutations in human disease. Nat Rev Genet, 6:389–402.

Tegunov, D. and Cramer, P. (2019). Real-time cryo-electron microscopy data preprocessing with Warp. Nat Methods, 16:1146–1152.

Teng, Y. et al. (2016). Mitochondrial ATAD3A combines with GRP78 to regulate the WASF3 metastasis-promoting protein. Oncogene, 35:333–343.

Tuppen, H. A. L., Blakely, E. L., Turnbull, D. M., and Taylor, R. W. (2010). Mitochondrial dna mutations and human disease. Biochimica et Biophysica Acta (BBA) - Bioenergetics, 1797:113–128.

Wiedemann, S. and Heckel, R. (2024). A deep learning method for simultaneous denoising and missing wedge reconstruction in cryogenic electron tomography. Nat Commun, 15: 8255.

Wu, L. J. and Errington, J. (2012). Nucleoid occlusion and bacterial cell division. Nat Rev Microbiol, 10:8–12.

Zhang, Y. et al. (2026). MISO regulates mitochondrial dynamics and mtDNA homeostasis by establishing membrane subdomains. Nat Cell Biol, 28:255–267.

Zhao, Y. et al. (2019). ATAD3A oligomerization causes neurodegeneration by coupling mitochondrial fragmentation and bioenergetics defects. Nat Commun, 10:1–20.

Zollo, C., Gomez Suarez, D., Bararpour, E. P., Jenner, A., Verma, P., Koch, J., Wilhelm, S., Jüngst, C., Faber, L., White, F., and García-Sáez, A. J. (2026). DRP1 and MID49 co-diffusion scans mitochondria for fission. Nature Cell Biology. Advance online publication.

