## Supplemental Figures for "ATAD3A structurally links mtDNA replication and mitochondrial fission"

### Supplementary Information

#### Figure S1

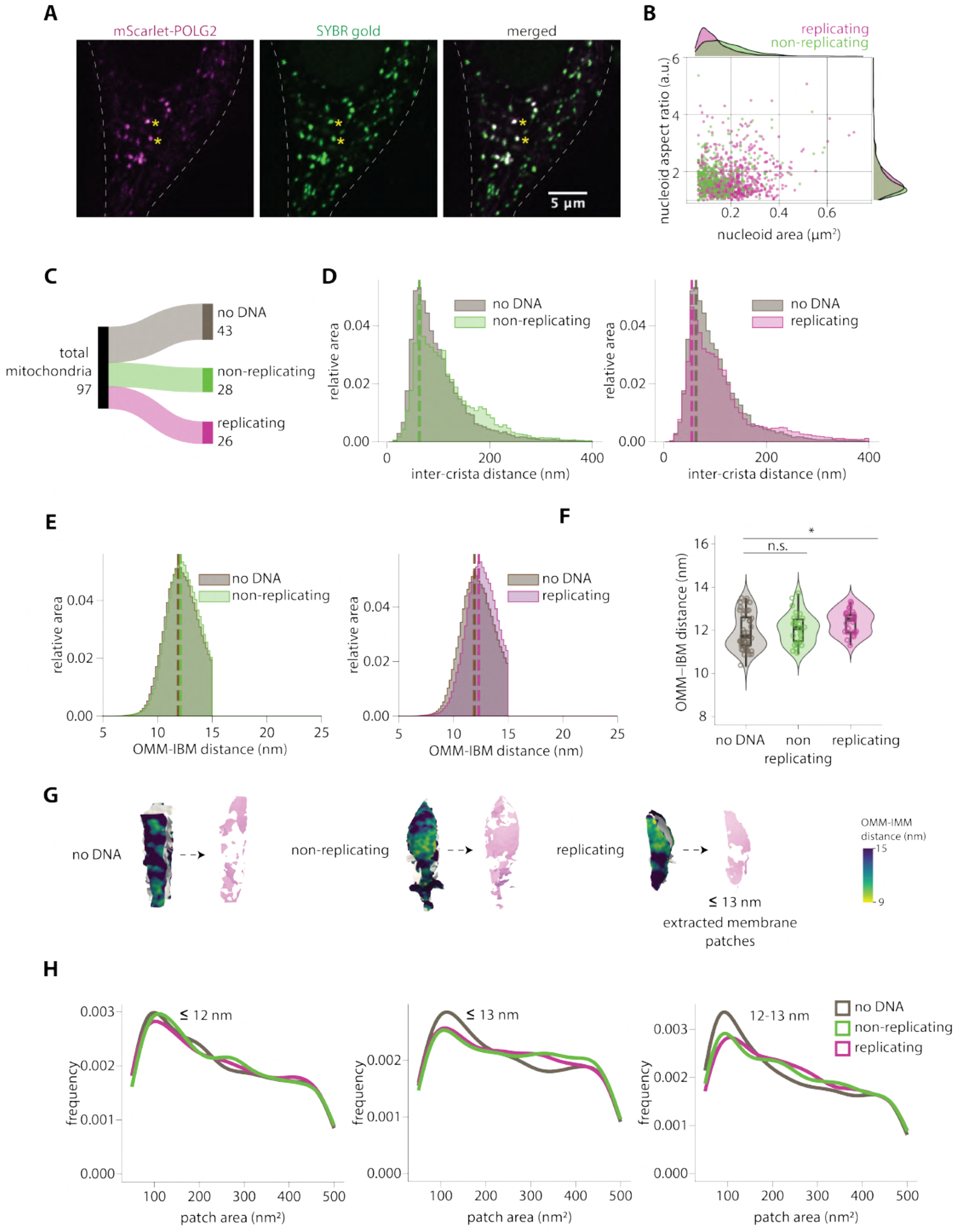

**Figure S1. mtDNA replication is associated with a unique membrane microenvironment.**
(**A**) Representative images of a cell expressing mScarlet-POLG2 and stained with SYBR Gold. Asterisks mark replicating mtDNA nucleoids. (**B**) 2D distribution of nucleoid area and nucleoid aspect ratio. Each dot represents a single nucleoid. (**C**) Classification of a total of 97 mitochondria into three classes based on fluorescence information. (**D**) Combined histogram of inter-crista distances for all three classes, dashed vertical lines correspond to peak histogram values of pooled data. (**E**) Combined histogram of OMM–IBM for all three classes, dashed vertical lines correspond to peak histogram values of pooled data. (**F**) Peak histogram values from each mitochondrion for OMM-IBM distance. Each dot represents peak value for the corresponding mitochondrion. (**G**) Representative examples showing extracted patches of OMM with different thresholds of OMM-IMM distance (example shown here for $\leq$13 nm). (**H**) Frequency distributions for area of patches extracted after applying different threshold for OMM-IMM distance. Number of mitochondria $= 43$ no DNA; 28 non-replicating, 26 replicating in D, E, F and H. Statistical significance in F is calculated using Mann-Whitney U test (***$p < 0.001$, **$p < 0.01$, *$p < 0.05$).

**Figure S2**

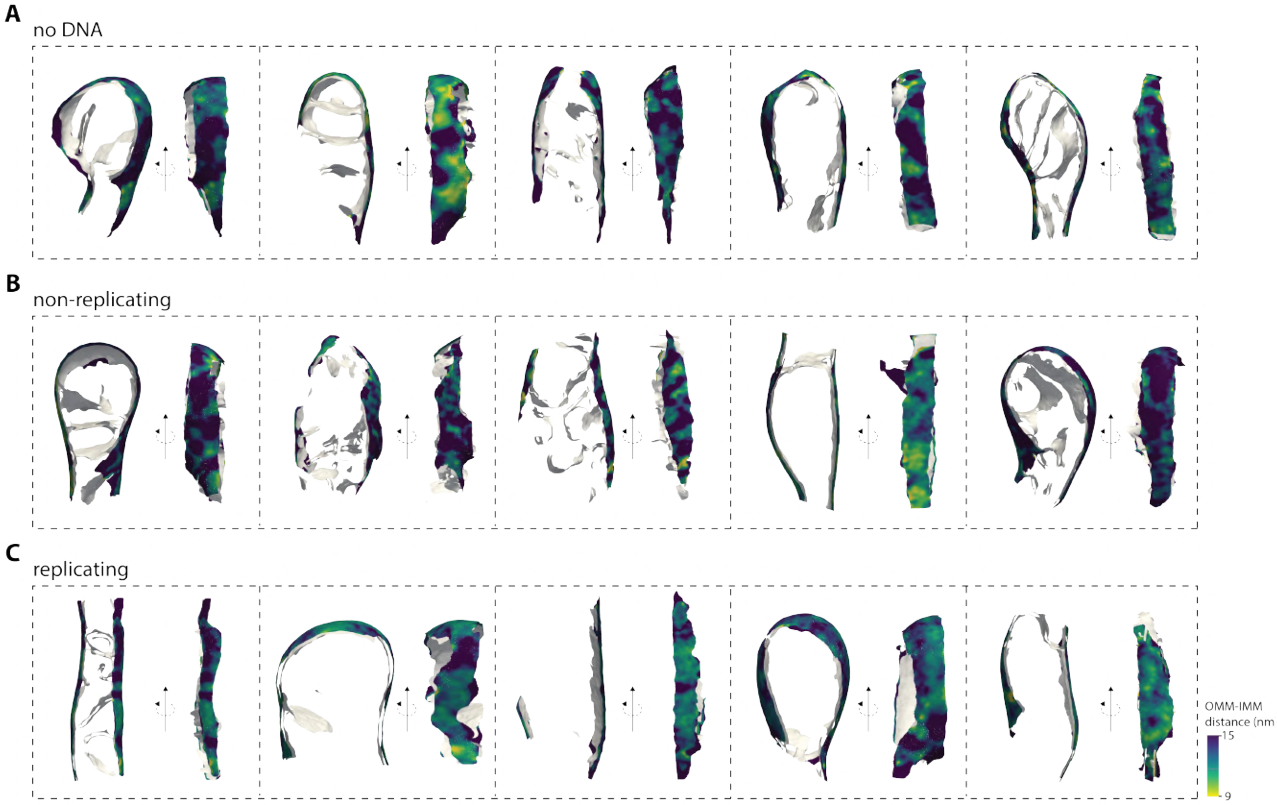

A
no DNA
B
non-replicating
C
replicating
OMM-IMM distance (nm
15
9

**Figure S2. mtDNA replication is associated with a unique membrane microenvironment.**
Gallery of representative mitochondria from (**A**) no DNA, (**B**) non-replicating, and (**C**) replicating categories with per-triangle OMM-IMM distance measurements mapped on the surface.

#### Figure S3

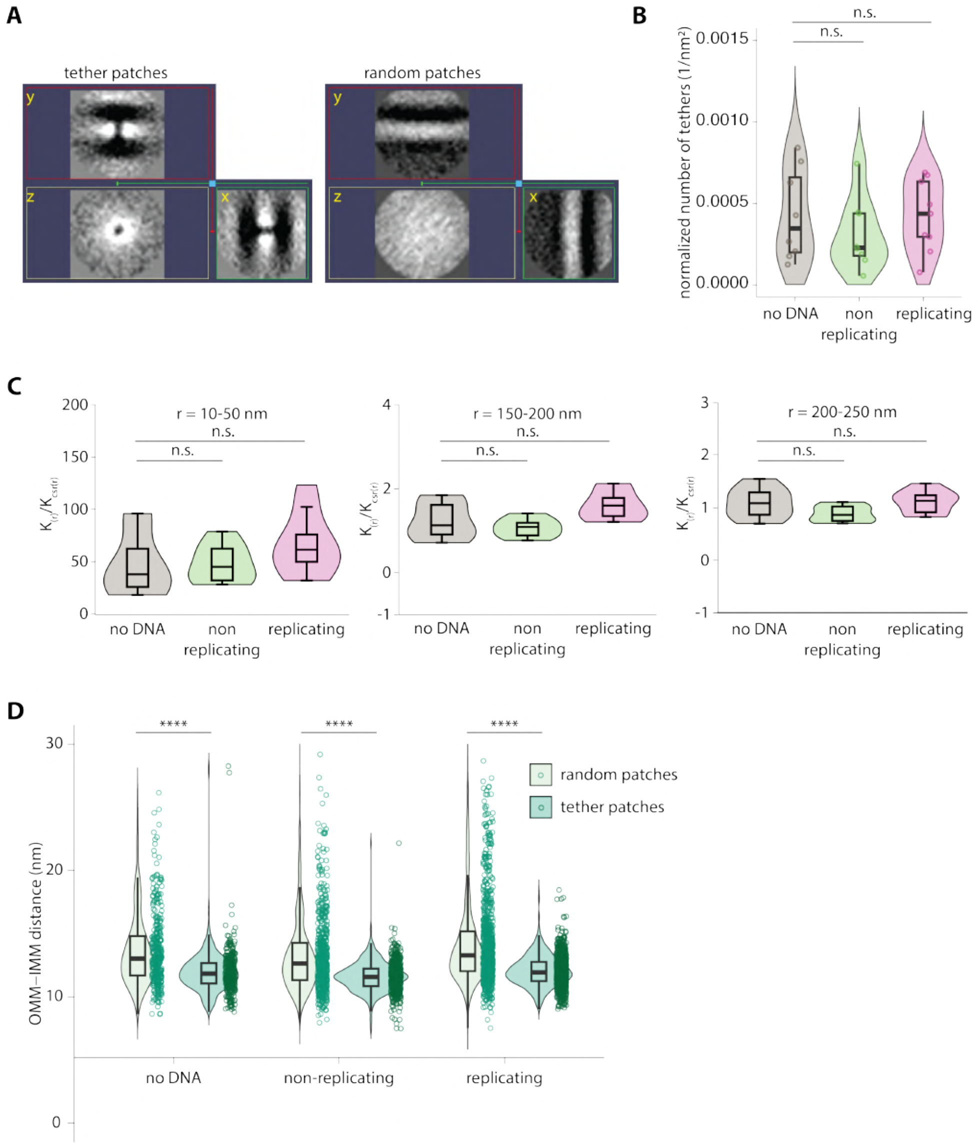
A
tether patches
random patches
B
normalized number of tethers (1/nm²)
0.0015
0.0010
0.0005
0.0000
n.s.
n.s.
no DNA
non replicating
replicating
C
r = 10-50 nm
r = 150-200 nm
r = 200-250 nm
n.s.
n.s.
no DNA
non replicating
replicating
D
OMM−IMM distance (nm)
****
random patches
tether patches
no DNA
non-replicating
replicating

**Figure S3. Regions of uniform OMM-IMM distance contain tether-like densities that are clustered at replicating mtDNA nucleoids.**
(**A**) X, Y and Z slice views of the global average from two-point manually picked tethers *(left)*; and X, Y and Z slice views of the global average from sub-tomograms extracted at random patches of the membrane *(right)*. (**B**) Number of tethers in each category of mitochondria normalized to OMM surface area are plotted as violin plot. Each data point represents one mitochondria. ($n = 8$ no DNA; 8 non-replicating, 9 replicating) (**C**) Quantification of the maximum value of $K(r)/K_{\mathrm{CSR}}(r)$ for each tomogram at the indicated radius intervals in each category. (**D**) OMM-IMM distance is plotted as a violin plot for random and tether containing patches of 10 nm radius within each category of mitochondria. Each data point represents median OMM-IMM distance for a patch. Statistical significance in B, C and D is calculated using Mann-Whitney U test (****$p < 0.0001$, ***$p < 0.001$, **$p < 0.01$, *$p < 0.05$)

#### Figure S4

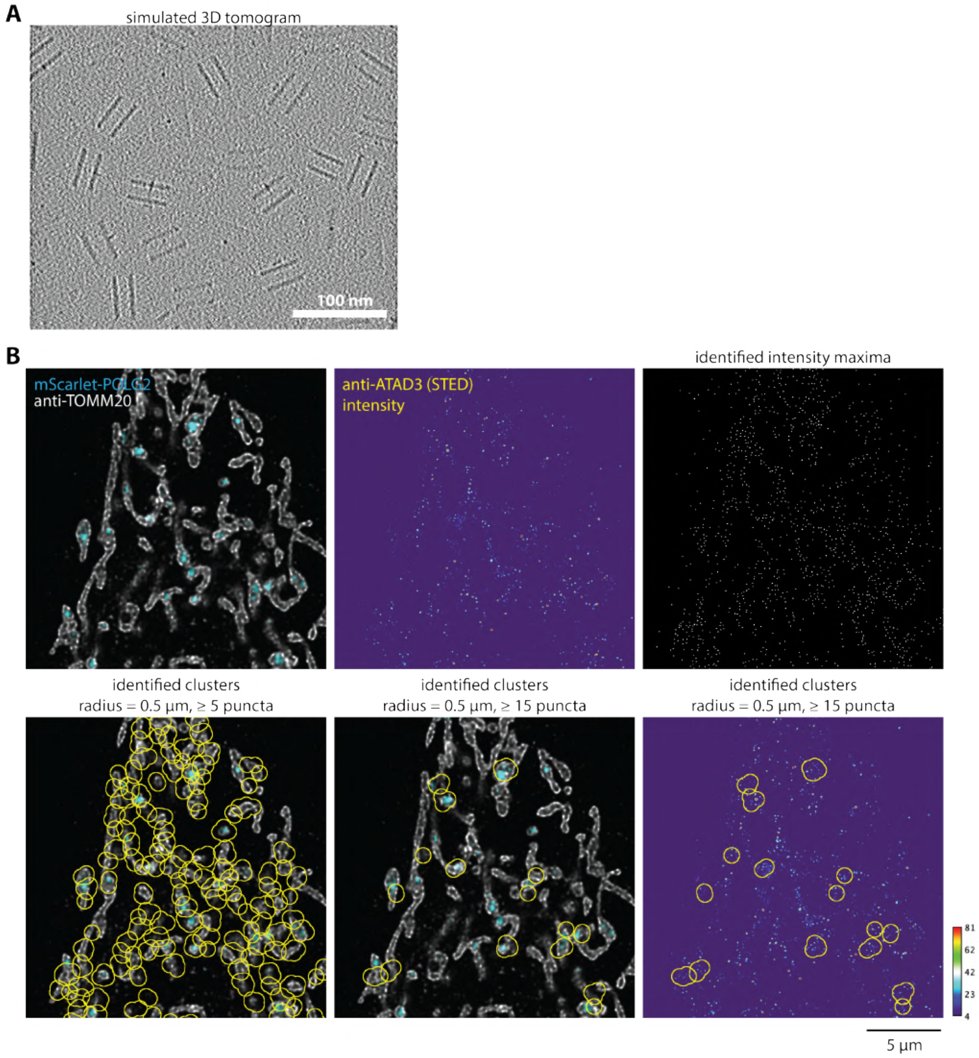
A
simulated 3D tomogram
100 nm
B
mScarlet-PGLC2
anti-TOMM20
anti-ATAD3 (STED)
intensity
identified intensity maxima
identified clusters
radius = 0.5 µm, ≥ 5 puncta
identified clusters
radius = 0.5 µm, ≥ 15 puncta
identified clusters
radius = 0.5 µm, ≥ 15 puncta
81
62
42
23
4
5 µm

**Figure S4. ATAD3A is enriched at replicating mtDNA nucleoids.**
(**A**) A simulated tomogram of the model generated using the AlphaFold model of ATAD3A hexamer with OMM and IMM PDB models. (**B**) Representative example showing STED resolved ATAD3A puncta. Each punctum is identified using the "Find Maxima" function in Fiji *(top)*. Clusters of ATAD3A puncta are identified with different thresholds. Each yellow ROI represents an identified cluster.

**Figure S5**

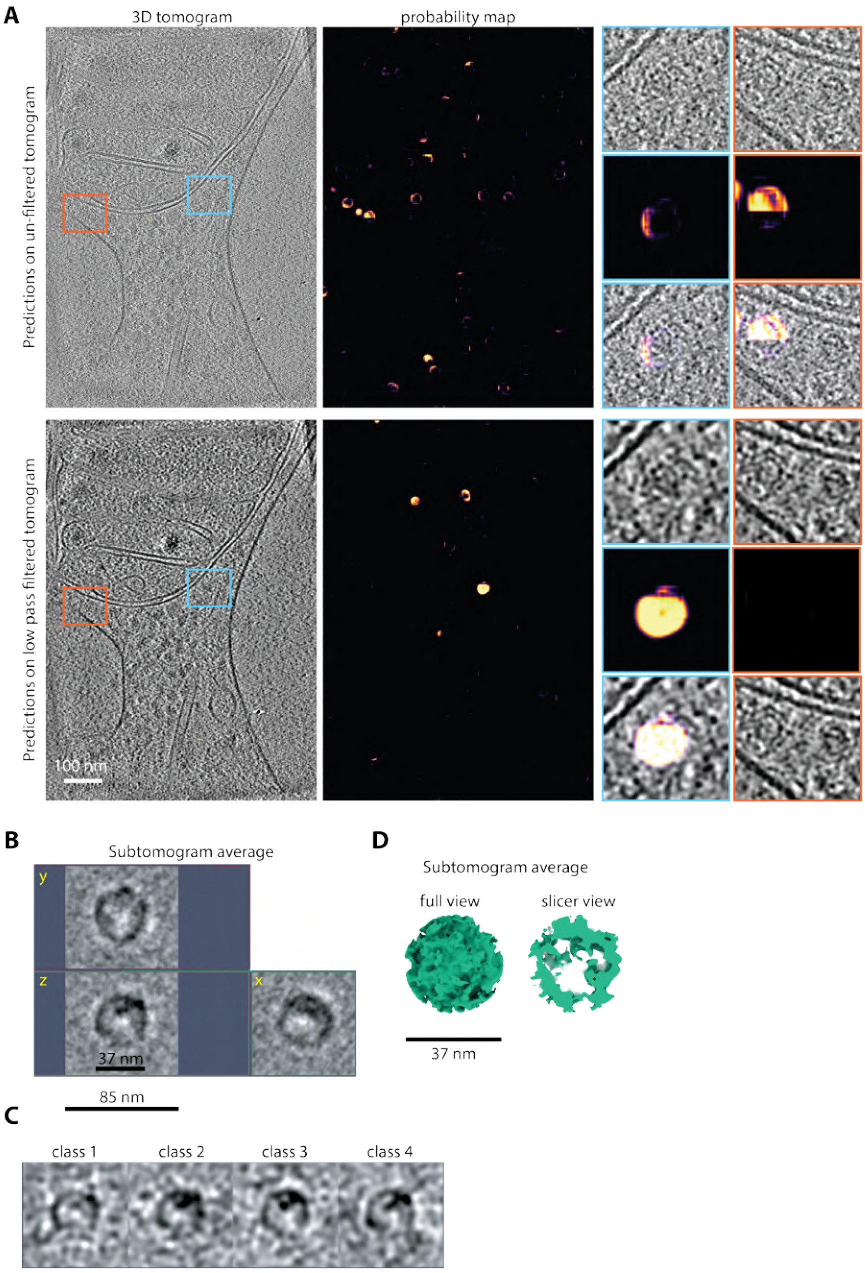
A
3D tomogram
probability map
Predictions on un-filtered tomogram
Predictions on low pass filtered tomogram
100 nm
B
Subtomogram average
y
z
x
37 nm
85 nm
C
class 1
class 2
class 3
class 4
D
Subtomogram average
full view
slicer view
37 nm

**Figure S5. GEM tags validate the identity of OMM-IMM ATAD3A tethers.**
(**A**) A z-slice from the 3D tomogram shown in Figure 4C with DeePiCt prediction probability maps for both the unfiltered tomogram and the low-pass filtered tomogram. Insets show examples of GEMs identified in both probability maps. (**B**) Subtomogram average of GEM particles identified using the DeePiCT model. (**C**) I3 classification of GEMs show 4 similar classes. (**D**) Slice view of the subtomogram average shows a hollow sphere of size ~37 nm.

**Figure S6**

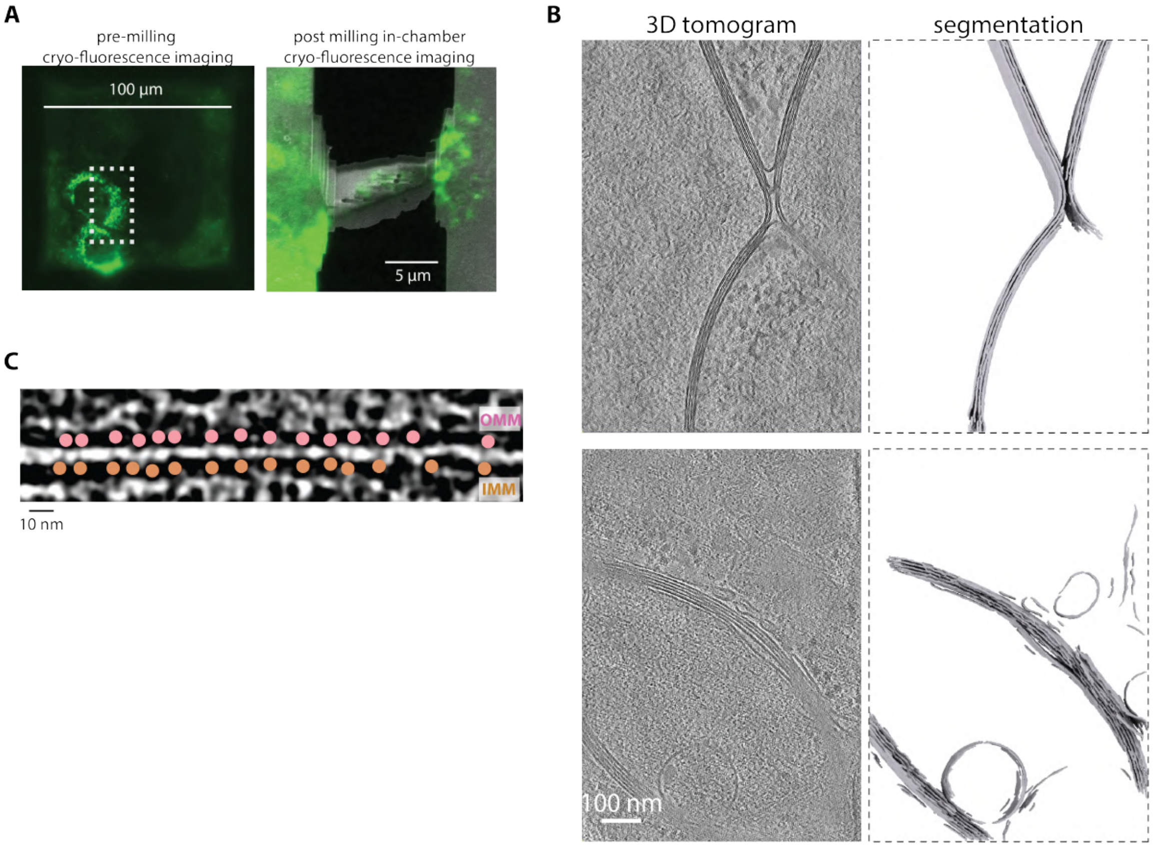
A
pre-milling
cryo-fluorescence imaging
100 µm
post milling in-chamber
cryo-fluorescence imaging
5 µm
B
3D tomogram
segmentation
100 nm
C
OMM
IMM
10 nm

**Figure S6. ATAD3A overexpression and knockdown remodels mitochondrial membranes.**
(**A**) Representative pre-milling cryo-fluorescence image of a cell overexpressing GFP-ATAD3A (ATAD3A *OE*, *left*). Post-milling cryo-fluorescence image correlated with SEM image, of a cell marked with a dotted rectangle (*right*). (**B**) Example 3D tomograms and membrane segmentations from ATAD3A *OE* cells, showing stacked mitochondrial membranes. (**C**) Inset showing uniform distance between OMM and IMM in ATAD3A *OE* cells with multiple tethering densities in between the membranes.

**Figure S7**

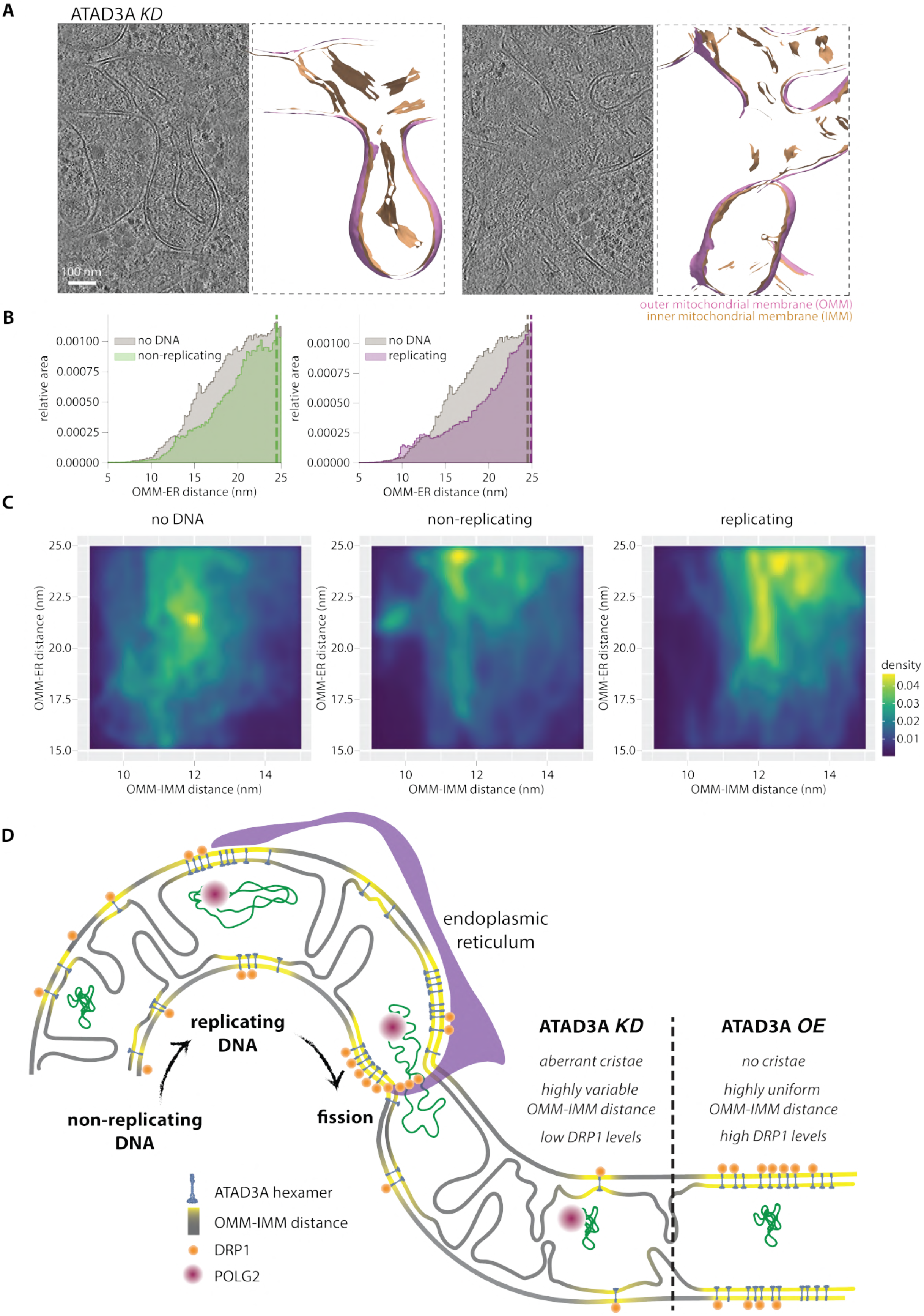

A
ATAD3A KD
100 nm
outer mitochondrial membrane (OMM)
inner mitochondrial membrane (IMM)
B
no DNA
non-replicating
replicating
relative area
OMM-ER distance (nm)
C
no DNA
non-replicating
replicating
OMM-ER distance (nm)
OMM-IMM distance (nm)
density
D
endoplasmic reticulum
replicating DNA
non-replicating DNA
fission
ATAD3A KD
aberrant cristae
highly variable OMM-IMM distance
low DRP1 levels
ATAD3A OE
no cristae
highly uniform OMM-IMM distance
high DRP1 levels
ATAD3A hexamer
OMM-IMM distance
DRP1
POLG2

**Figure S7. ATAD3A overexpression and knockdown affect recruitment of DRP1 and mitochondrial morphology.**
(**A**) Representative example slices from tomograms collected on ATAD3A *KD* cells showing branched mitochondria. (**B**) Combined histogram of ER-OMM distances for all three classes, dashed vertical lines correspond to peak histogram values of pooled data. (**C**) Heatmap for 2D distribution of ER-OMM vs OMM-IMM distance for each triangle for all three classes. Statistical significance in A is calculated using unpaired $t$-test (****$p < 0.0001$, ***$p < 0.001$, **$p < 0.01$, *$p < 0.05$). (**D**) Model showcasing differences across different states of mtDNA replication in ATAD3A levels and corresponding changes in OMM-IMM distance and recruitment of the mitochondrial fission machinery (DRP1 and endoplasmic reticulum). Consequences of ATAD3A knockdown and ATAD3A overexpression on cristae architecture, OMM-IMM distance, and DRP1 levels are displayed.
